# Histone H3K27 methyltransferase EZH2 interacts with MEG3-lncRNA to directly regulate integrin signaling and endothelial cell function

**DOI:** 10.1101/2022.05.20.492787

**Authors:** Tatiana Dudnakova, Hywel Dunn-Davies, Antonella Nogara, Julie Rodor, Anita Thomas, Elisa Parish, Philippe Gautier, Alison Meynert, Paolo Madeddu, Andrea Caporali, Andrew Baker, David Tollervey, Tijana Mitić

**Affiliations:** University and British Heart Foundation Centre for Cardiovascular Science, Queen’s Medical Research Institute, The University of Edinburgh, 47 Little France Crescent, Edinburgh, EH16 4TJ, UK; Wellcome Centre for Cell Biology, University of Edinburgh, Michael Swann Building Max Born Crescent, King’s Buildings, Edinburgh EH9 3BF, UK; Bristol Medical School, Translational Health Sciences, University of Bristol, Research and Teaching Floor Level 7, Queens Building, Bristol Royal Infirmary, Bristol BS2 8HW, UK; MRC Human Genetics Unit, MRC Institute of Genetics and Cancer, The University of Edinburgh, Western General Hospital, Crewe Road, Edinburgh EH4 2XU, UK

## Abstract

Enhancer of Zeste Homologue 2 (EZH2) modulates gene transcription during endothelial cell (EC) dysfunction, via interaction with non-coding RNAs (ncRNAs). Thus, EZH2 can act as a rheostat in deposition of histone H3K27 trimethylation (H3K27me3) to repress many genes. We profiled EZH2-RNA interactions using <u>f</u>ormaldehyde/UV assisted cross-linking <u>l</u>igation <u>a</u>nd <u>s</u>equencing of <u>h</u>ybrids (FLASH-seq) in primary human ECs. Transcriptome-wide EZH2-associated ncRNAs and RNA–RNA interactome were obtained. This approach revealed EZH2 directly binding maternally expressed gene (MEG3) and MEG3:MEG3 hybrid structures. By chromatin immunoprecipitation with sequencing (ChIP-seq) following depletion of MEG3, we discovered that MEG3 targets and controls recruitment of EZH2/H3K27me3 onto a regulatory region of integrin subunit alpha 4 (ITGA4). MEG3 knockdown or pharmacological inhibition of EZH2 de-repressed *ITGA4,* whilst improving endothelial cell function *in vitro,* and increasing *ITGA4* expression *in vivo*. Our study demonstrates new role for MEG3, as instrumental in epigenetic regulation of EC function by EZH2, through targeting of integrin-dependent signalling.

## INTRODUCTION

The intact vascular endothelium is essential for maintaining vascular homeostasis, providing endothelial cells (ECs) with a barrier to external stimuli in health and disease. Changes in endothelial function in response to stimuli impose changes in mechanical properties, involving cytoskeletal rearrangements critical for ECs morphogenesis. Multiple extracellular matrix (ECM) proteins and cell surface ligands are involved in all types of cellular adhesion, as mediators of the interactions between ECs and intracellular cytoskeleton [1]. The detailed mechanisms behind these processes remain largely unexplored, but the role of non-coding RNA (ncRNAs) in directing epigenetic changes in ECs is being more recognised. The long non-coding RNA (lncRNAs) can interact with chromatin and recruit protein complexes to remodel chromatin states and alter the epigenetic landscape, thus regulating transcription and gene expression [2]. Several nucleus-localising lncRNAs have been reported to interact with the transcriptional co-repressor, Polycomb Repressive Complex 2 (PRC2) [3]. These interactions appear to be cell-type specific and context dependent, yet only <2% of lncRNAs have been experimentally characterised [4]. The PRC2 comprises core subunits, EED, EZH1/2, SUZ12, and RBBP4/7, all are required to guide the enzymatic activity of EZH2 in depositing histone H3 lysine K27 tri-methylation (H3K27me3) repressive mark [5]. The H3K27me3 mark is implicated in numerous cardiovascular diseases involving ncRNA and endothelial dysfunction, and it strongly represses key cardiac genes [6].

In this study, we investigated the direct binding of RNAs to EZH2 (catalytic component) in ECs. A significant proportion of EZH2-associated RNA transcripts were obtained. Using a hybrid detection approach, we also examined functional interaction between EZH2 and RNA species in primary ECs. We showed that the RNA binding capacity of EZH2 was relative to its ability to capture the RNA-RNA interactions (hybrid reads); adding to its prevailing chromatin binding capacity.

The large nuclear retained lncRNA, MEG3 (<u>m</u>aternally <u>e</u>xpressed <u>g</u>ene 3) is located on chromosome *14q32.3* in humans, and was shown to associate with chromatin [7]. Increased MEG3 expression in primary ECs is both age-dependent and hypoxia-induced (e.g. during ischaemia), and a contributor to pulmonary hypertension [8, 9]. MEG3 antagonises angiogenesis, as increased brain vasculature was observed in the MEG3 knockout embryos, further suggesting pro-angiogenic role of MEG3 inhibition [10, 11]. Depletion of MEG3 improved angiogenic responses following femoral artery ligation (limb ischaemia) in aged mice and attenuated endoplasmic reticulum stress-mediated apoptosis following myocardial infarction, both leading to blood flow recovery [12]. In all, MEG3 is a valid target to improve postischemic angiogenesis. Moreover, MEG3 regulates vital pathways in EC biology, by enhancing stability and transcriptional activity of p53 and targeting TGF-β [13]. These effects are believed to occur via lncRNA-guiding the deposition of EZH2 transcriptional repressor, akin to its blocking of target genes in cancer cells and pluripotent stem cells, which typically express MEG3 at a low level, unlike primary cells [14]. The removal of H3K27me3, through PRC2 inhibition, increases recovery post limb ischaemia [15] and improves aortic performance in a model of thoracic aortic aneurysm [16]. Similar events are seen in other disease states known to arise from endothelial dysfunction [17, 18], however the involvement of MEG3 in guiding EZH2 actions in the dynamic processes of endothelial dysfunction has not been studied yet.

Hence, we hypothesize that EZH2 directly interacts with MEG3 and has potential functional importance in repressing vascular genes expression. Here we show that many types of RNA transcripts were directly associated with EZH2, and that EZH2 specifically targets MEG3:MEG3 structures in primary ECs. Next, we identified and validated MEG3-targets. Further mechanistic investigation showed that MEG3 was directly involved in guiding EZH2 and depositing H3K27me3 methylation on the regulatory region of integrin gene, *ITGA4*. We established that the MEG3-EZH2 complex represses EC migration and adhesion to fibronectin, by tempering integrin receptor signaling to facilitate interactions with ECM. In all, we have demonstrated that MEG3 is instrumental for epigenetic regulation of EC function by PRC2, through targeting the integrin-dependent signalling.

## RESULTS

### EZH2-FLASH identifies direct endothelial RNA targets

To unravel the interactions of EZH2 with RNA (EZH2-RNA interactome), we performed formaldehyde and UV cross-linking assisted ligation <u>a</u>nd <u>s</u>equencing of <u>h</u>ybrids (FLASH) [19–21] (**Fig. 1a**). The use of FLASH approach has specifically allowed us to identify ***1***- direct RNA targets (*single reads*) and ***2-*** RNA-RNA interactions (*hybrid reads*) associated with the candidate protein, i.e. EZH2. To precipitate EZH2, we used anti-EZH2 antibody and crosslinked cell lysates of human umbilical vein ECs (HUVECs). We used non-immune (mouse or rabbit) IgG as a control. RNA bound to EZH2 was isolated, followed by deep sequencing and data analysis, to obtain endothelial transcriptome with given RNA biotypes (**Fig. 1b**). Comparable data were obtained from two independent biological replicates **(Supplementary Table S1)** confirming that FLASH, with its stringent purification steps, is a robust method to identify endothelial RNA species and base-pairing interactions with EZH2. The differential enrichment (DEseq2) for each EZH2-associated transcript was determined compared with IgG control pulldown [22]. Statistically significant differences were identified in >15,000 peaks (N1: 15,399 and N2: 16,768) and a list of significant differentially expressed EZH2-FLASH targets; with RNA gene classification, i.e. biotypes (log_2_FC>0.5) is given in **Supplementary Table S1**.

**Figure 1.**
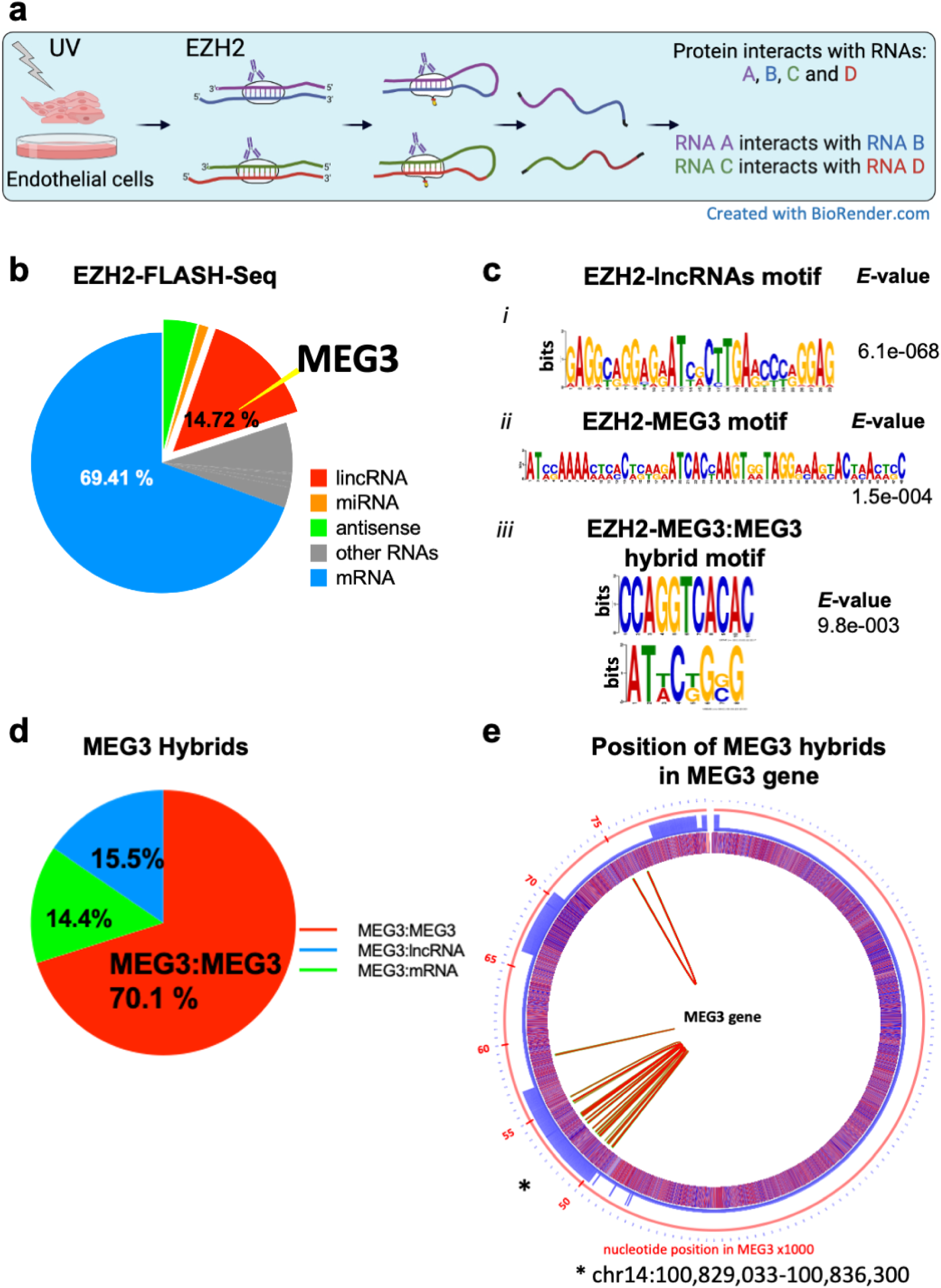
**a.** Schematic representation of steps in FLASH-seq (formaldehyde and UV cross-linking, ligation, <u>a</u>nd <u>s</u>equencing of <u>h</u>ybrids) with EZH2 immunoprecipitation using lysates from UV crosslinked endothelial cells. Dynamic EZH2-RNA complex formation occurs as represented. Following RNA ligation and chimera formation between interacting RNAs, sequencing is performed. Further analysis of single and hybrid reads bound by EZH2, reveals interacting RNA molecules. **b.** Distribution of annotated reads over genome, with gene classification (biotype), from statistically filtered EZH2-FLASH data with two biological replicates in HUVECs and MEG3-lncRNA (yellow wedge) as the candidate. **c. *I and ii*** Enriched motifs with sequences in MEG3 mRNA of EZH2-FLASH that uniquely overlap exons; the logos were drawn using the top 4-8nucleotides K-mers for each experimental replicate (***top*** and ***middle***) and z-score for each. Motif analysis was performed using the MEME suite (Bailey et al., 2009) [33] ***iii***: Enriched motif within the fragments of MEG3:MEG3 hybrids **d.** Total RNA-RNA interactions associated with MEG3 at chr14:101292445-101327360, MEG3 id = **NR_002766.2**) and distribution of all MEG3 interactions among various classes of RNAs as captured by EZH2-FLASH. **e.** Intermolecular MEG3-RNA interactions found in chimeras captured by EZH2-FLASH. Chimera counts were mapped for all genomic features of annotated hybrids and the ones of MEG3 were plotted in the circos plot with position along the MEG3 genomic sequence. The main MEG3 hybrid is MEG3 and are represented by the number of interactions in red. The feature as a line: *Red* circle shows the position in the MEG3 gene in kilobases with * 50-55kb falling within exon3; *Blue* circle is a visual representation of MEG3 exons. Regions overlapping exons are represented in solid blue. *Purple* broad circle shows the nucleotides. The nucleotides at each position are: **A**: dark blue, **C**: light blue, **T**: light red, **G**: dark red. The details on the feature: The inner part of the white circle shows MEG3:MEG3 hybrids; Arcs connecting the centre of each hybrid fragment are shown in red, and the regions spanned by the hybrid fragments are shown in light green.

EZH2/PRC2 has been reported to preferentially target nascent transcripts [23]. Our data is consistent with this, with 30% of single FLASH reads mapped within intronic and intergenic regions represented as other RNAs (in gray, **Fig. 1b**), indicating that *bona fide* interactions are recovered. We identified total of 390 (log_2_FC>0.5) novel lncRNAs and found known lncRNA targets. Amongst the top 20 represented lncRNAs, bound to EZH2, *MEG3* gene (ENSG00000214548.14) was recovered (**Supplementary Fig. 1A; Supplementary Table S1)** with single and hybrid hits detected in both replicates **(Supplementary Table S2** and **S3,** respectively). Resultant sequences of EZH2 single hits have been validated and they aligned with MEG3 gene sequence, as compared by Pairwise Sequence Alignment (https://www.ebi.ac.uk/Tools/msa/clustalo/) [24]. We further focused on MEG3-lncRNA for analysis. High EZH2 binding was seen over exons regions of MEG3 RNA, supporting functional interactions (∼75% over *MEG3* exons 3-7 and 25% on introns) (**Supplementary Fig. 1B**). More than one annotated transcript was derived from the *MEG3* genomic region, and most EZH2 reads were distributed over ENST00000451743.6 (**NR_002766.2,** transcript variant 1) relative to the ENST00000423456.5 transcript (**NR_003530,** transcript variant 2**; Supplementary Fig. 1C),** as per Ensembl release 77 [25]. The MEG3 ENST00000451743.6 transcript is already known to be highly expressed in aorta and tibial artery according to GTEx profile https://gtexportal.org/home/transcriptPage. In a separate RNA immunoprecipitation experiments we precipitated the repressive chromatin (EZH2 and H3K27me3 mark, see methods) using UV-crosslinked lysates of ECs and verified interaction with MEG3. Following RNA isolation, qPCR analyses detected MEG3 with maximum recovery over exon 3, against which primers were designed (**Supplementary Fig. 1D**). To a lesser extent the binding was also observed for other PRC2 subunits, SUZ12 and EED. Motif analysis using MEME identified enriched sequence motifs within MEG3 fragments associated with EZH2 (**Fig. 1c**). These motifs comprised “GTGA” and “AGGA” nucleotides. Similar motifs were identified for all other endothelial RNAs detected in our work and have been observed by others before for EZH2-bound lncRNAs in a different cell type [3].

### EZH2 targets structures within MEG3

RNA-RNA duplexes associated with EZH2 as chimeric cDNA species were identified in two independent FLASH experiments (biological replicates; 3,944 hybrids from N1 and 3,228 hybrids from N2). These revealed the predominance of MEG3:MEG3 base-pairing (**Fig. 1d**) over MEG3 base-pairing with other RNAs. In addition, specific MEG3:MEG3 base-pairing sites were identified in the replicate datasets (**Supplementary Fig. 1E**) and are representative of MEG3 intermolecular secondary structure. The stability of the predicted base-pairing between the two RNAs in the chimeras was calculated as the change in free energy of hybridization (ΔG in kcal mol^-1^ < -10) and 553 interactions passed these filters. Hybrids counts were plotted against the predicted binding energy of MEG3 interactions for EZH2-FLASH ***(i)*** and IgG-FLASH (***ii***, **Supplementary Fig. 1F *i*** and ***ii***). Chimera counts were mapped for all genomic features of annotated hybrids and represented with their position along the MEG3 genomic sequence. “*Viennad*” files for clustering of MEG3:MEG3 interactions (hybrids) were generated identifying the nucleotides that interact by base pairing and giving mostly likely base pairings form with *dG* energy of folding predictions (**Supplementary Table S3**). The representation of MEG3:MEG3 interactions found in chimeras captured by EZH2-FLASH are seen in circos plot (**Fig. 1e**). The majority of these interactions were between MEG3 positions 52623-52670, corresponding to highly conserved exon 3, and positions 75277-76613.

### MEG3 directly targets angiogenic genes

To unravel interactions and functions of chromatin associated *MEG3* lncRNA in ECs we identified genomic binding sites by obtaining DNA targets using chromatin isolation by RNA purification (ChIRP). Published biotin-labelled antisense probes against MEG3 were used to target MEG3:RNA and MEG3:DNA complexes, that were then captured with streptavidin magnetic beads (**Supplementary Fig. 3A**) [26]. The specificity of biotin-labelled probes for MEG3 gene, as opposed to other genes, was confirmed using one-step qPCR with RNA purified from ChIRP (**Supplementary Fig. 3B**). We performed chromatin isolation after glutaraldehyde crosslinking, followed by RNA purification for qPCR and DNA purification for sequencing. *MEG3* ChIRP-Seq identified >5000 peaks (**Fig. 2a**) and motif enrichment analysis (MEA) of regulatory sequences, revealed top two consensus DNA motifs given in **Fig. 2b**. Computational analysis with pipeline for ChIRP-seq data processing is outlined in **Supplementary Fig. 3C.** Identified genomic sites of MEG3 interactions were considered as true MEG3-targets if they were recognized in both independent ChIRP experiments, and undetected in the negative control (using probes against *LacZ* RNA). We detected 4742 and 3275 MEG3-binding loci and true peaks identified from two replicates, of which 863 and 982 (respectively) were associated with genes or gene regions and showed peak heights ≥3; 347 of these were mRNA targets common to both experiments, suggesting that they are indeed MEG3 targets (**Supplementary Table S4)**. Identified MEG3 genomic loci were mostly associated with the intergenic and intronic regions, as well as promoter regions. The presence of many putative distant binding sites away from the promoter, for endogenous *MEG3* lncRNA has further suggested an enhancer regulation, which we examined too. Through assessing the genomic regions of MEG3 occupancy in HUVECs we overlapped the regions with novel and known enhancers and then attributed these loci to the nearest genes via their functional pathways (identified ontologies in **Table 2**). The total list of identified enhancers and super enhancers within the MEG3-associated genomic loci with nearby genes is given in **Supplementary Table S5**.

**Figure 2.**
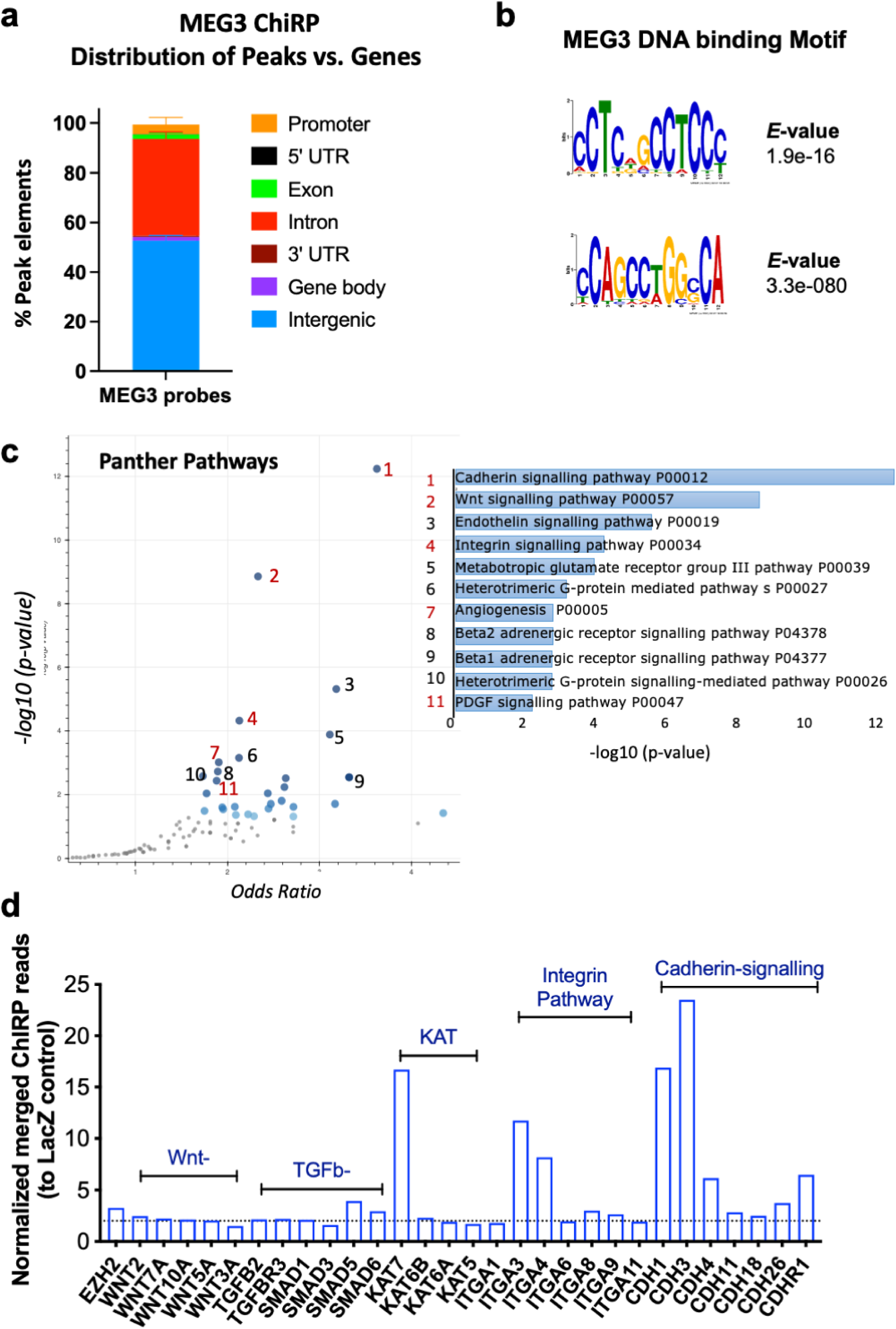
***a.*** ChIRP-seq analysis showing % of peaks elements in the pull down using biotinylated probes against MEG3 gene. The ChIRP pull-down with LacZ oligoes was used as negative control. *N*=2 biological replicates followed by bioinformatics analysis to merge two replicates to boost the signals. ***b.*** Top enriched motifs by E-value (statistical significance as calculated by MEME) within the DNA sequence that uniquely overlapped regions associated with MEG3 probes. Motif analysis was performed using the MEME suite (Bailey et al., 2009) [45]. ***c.*** Volcano plot of gene enrichment analysis for all MEG3 peaks-associated genes (**left**) with top rated GO biological pathway annotations (**right**). Enrichr analysis with Panther pathway resource was used to associate most represented genes with pathways (Mi, H and Thomas P, 2009) and *p*-Value was calculated by the Binomial statistic with cutoff of 0.05 used as a start point. ***d.*** MEG3 binding to genomic loci in HUVECs. Pull downs for ChIRP-seq were performed with biotinylated MEG3-probes or control LacZ probes. Merged reads were related to control LacZ reads and expressed as enrichment. Background level was defined by LacZ negative control probes in merged ChIRP-seq and enrichment ≥3 was considered significant. Relevant genes were plotted from the most represented pathways of Cadherin and Integrin signalling and lysine acetyl transferase (KAT).

**Table 1.**
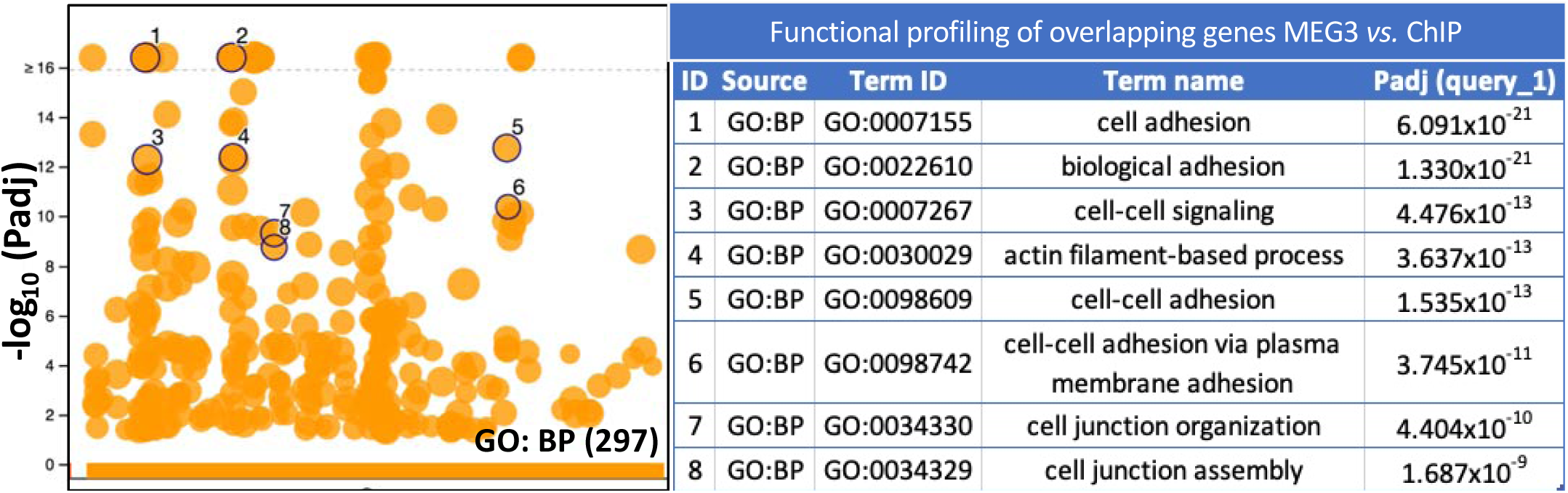
Profile (left) with tabulated named pathways (right) for top enriched mRNA targets found as per **Fig. 3c**. Maximum peak score of ChIP signal for EZH2 and H3K27me3 intersected with top enriched MEG3 peaks generating list of common genes that were functionally profiled. Overlap of the number of genes mostly belong to the biological pathway regulating cell adhesion.

**Table 2.** Gene enrichment analysis with functional profiling of top identified peaks associated with enhancers in MEG3 genomic targets. The enhancer regions were associated with nearest genes and sorted based on their linked pathway. The total summary of identified enhancers, their chromosomal location, associated super-enhancer and nearest genes is deposited as **Supplementary Table S5.csv file**

| Functional profiling of identified enhancer-associated MEG3-genes |  |  |  |  |
| --- | --- | --- | --- | --- |
| ID | Source | Term ID | Term Name | padj (query_1) |
| 1 | GO:BP | GO:0007015 | actin filament organization | $7.558 \times 10^{-4}$ |
| 2 | GO:BP | GO:0032970 | regulation of actin filament-based process | $2.503 \times 10^{-3}$ |
| 3 | GO:BP | GO:0030029 | actin filament-based process | $2.503 \times 10^{-3}$ |
| 4 | GO:BP | GO:0032956 | regulation of actin cytoskeleton organization | $7.299 \times 10^{-3}$ |
| 5 | GO:BP | GO:0110053 | regulation of actin filament organization | $1.958 \times 10^{-2}$ |
| 6 | GO:BP | GO:0150116 | regulation of cell-substrate junction organization | $2.794 \times 10^{-2}$ |
| 7 | GO:BP | GO:0042060 | wound healing | $2.992 \times 10^{-2}$ |
| 8 | GO:BP | GO:0007010 | cytoskeleton organization | $3.825 \times 10^{-2}$ |
| 9 | GO:BP | GO:0051493 | regulation of cytoskeleton organization | $2.794 \times 10^{-2}$ |
| 10 | GO:BP | GO:0045944 | positive regulation of transcription by RNA polymerase II | $2.387 \times 10^{-2}$ |

The function of MEG3-associated genes was assigned using Panther Pathway and Enrichr classifications to identify pathways (**Supplementary Table S4**) with direct relevance to endothelial repair processes [27]. The occupancy was seen on multiple genes belonging to Cadherin signalling (*CDH1* and *CDH3*), Wnt signalling (Wnt, beta-catenin) and Integrin (*ITGA3*, *ITGA4*) signalling, all related to the process of angiogenesis *P00005* (**Fig. 2c**). Further enrichment was found for groups of genes annotated with Gene Ontology (GO) terms related to *cell adhesion* (GO:0007155), focal adhesion and as well as cell adhesion via plasma membrane (GO:0098742). Normalized ChIRP peak reads over named genes are given as enrichment in **Fig. 2d**. Amongst the MEG3 targets we found the genomic locus of *EZH2* (chr7:148,806,055-148,900,311), signifying *MEG3* physical interaction with the *EZH2* gene (**Supplementary Fig. 3D)**. However, all peaks belonging to TGF beta pathway, although detected, were at the background level (peak for TGFBR3=3) apart form SMAD5 and SMAD6. The qPCR validation of *EZH2* and other MEG3-genomic targets was performed in separate ChIRP experiments. Enrichment of MEG3 signal by ChIRP-qPCR versus negative control over selected targets is seen in **Supplementary Fig. 3E**.

### MEG3 genomic target sites in ECs overlap with chromatin occupancy by EZH2 and H3K27me3

To study the relationship between MEG3-regulated targets and EZH2-chromatin binding sites, in human ECs, we next performed bed intersection of ChIRP-seq files with those from public ChIP-seq. H3K27me3 chromatin marks are recognised by the PRC2 core subunit EED to promote PRC2 engagement. The enrichment profile of H3K27me3 can depend on the cellular context, hence we systematically analysed the genome-wide profile of H3K27me3 tracks from the Gene Expression Omnibus (GEO) datasets of ChIP-seq performed in HUVECs. We searched for the GEO datasets in the NCBI portal (ncbi.nlm.nih.gov) using key words of “EZH2 or H3K27me3 ChIP-seq in HUVECs”. The following datasets were found, for EZH2 ChIP under accession number <u>GSE109626</u>; and <u>GSM733688</u> and <u>GSM945180</u> for H3K27me3 ChIP. We next sought to identify peaks within the MEG3-ChIRP-seq data that overlapped with the peaks for chromatin occupancy by EZH2 or H3K27me3; as well as to map targeted enhancer regions (**Fig. 3a**). The ChIRP and ChIP signal convergence per gene region and type of genes are represented in **Fig. 3b**. We functionally profiled the overlapping mRNAs with their molecular function and biological processes. All overlapping targets are listed in **Supplementary Table S6**. We have also mapped the enhancer regions occupied by MEG3 peaks as reported in **Supplementary Table S5**.

**Figure 3.**
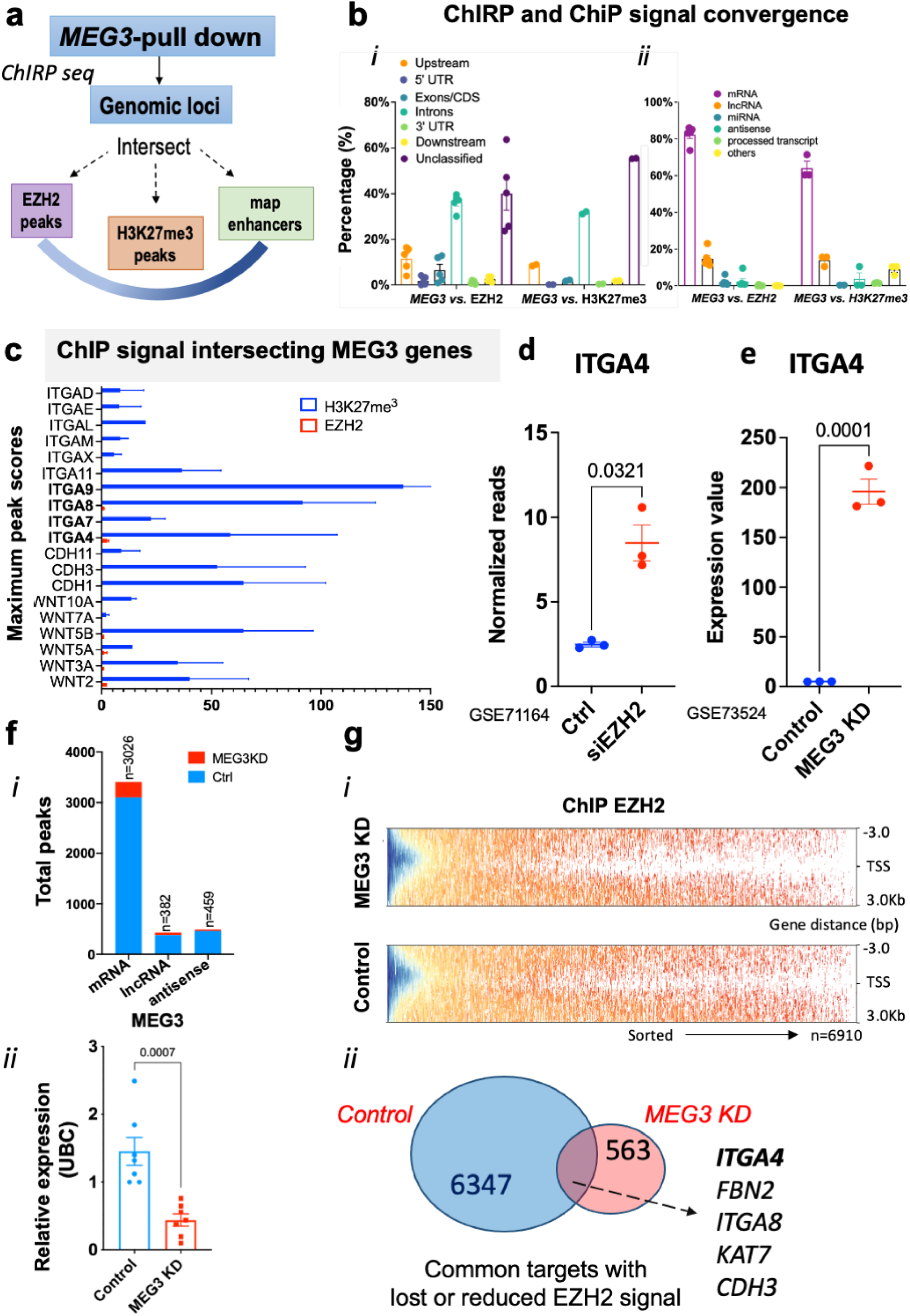
**a.** Overview of the critical steps to obtain MEG3-bound genomic loci and intersections with EZH2 and H3K27me3 signals (obtained from GEO databases for HUVECs). In addition, enhancer regions were mapped within the genomic tracks. The intersection between GEO EZH2 ChIP data, GEO H3K27me3 ChIP data and statistically filtered MEG3-ChIRP data from two biological replicates was performed. The number of genes and degree of overlap is obtained between MEG3 and PRC2-dependent genes. The *p-values* are a result of hypergeometric test. **b.** Distribution of MEG3 peaks overlapping EZH2-ChIP peaks or H3K27me3-peaks with intersecting reads in relation to ***(i)*** gene regions and ***(ii)*** gene-type. **c.** Maximum peak score of ChIP signal for EZH2 and H3K27me3 intersecting the top enriched MEG3 peaks associated with nearest genes. Highest EZH2 peak score is over ITGA4, whereas H3K27me3 was detected in ITGA4, ITGA7, ITGA8 and ITGA9, members of ITGA family. **d.** Normalized reads from RNA-seq de novo analysis of GEO: GSE71164 dataset on Hg38, and expression of ITGA4 gene between Scr and siEZH2 depleted HUVECs, showing that ITGA4 is targeted by EZH2. Dataset in *d* and *e* is compared using Student’s t-test. **e.** ITGA4 expression from microarray analysis in C2C12 cells depleted of MEG3 (10nM, LNA GapMer) as per GEO dataset: GSE73524. The data shows that ITGA4 is a direct target of MEG3. **f. *(i)*** Total number of representable peaks (mRNA, antisense and lncRNA genes) from ChIP-seq analysis of Scr *vs.* MEG3 KD HUVECs. ***(ii*)** Depletion of MEG3 gene in HUVECs (10nM LNA gapmers) was achieved with relative expression showing ∼70% reduction compared with Scr control. **g. *(i)*** Heat map showing distribution of reads and EZH2 densities at all unique RefSeq genes within TSSs ± 3 kb, sorted by EZH2 occupancy, in Control *vs.* MEG3 deficient (10nM) HUVECs. ***(ii)*** Overlap of ChIP-results between MEG3 and EZH2-dependent genes, with overlapped genes belonging to the biological pathway regulating cell adhesion. The common targets had lost or reduced EZH2 ChIP-signal.

We performed the overlap between the MEG3 binding sites and the H3K27me3 and EZH2 occupancy, respectively. Computational analysis with pipeline is given in **Supplementary Fig. 4**. The ChIP enrichment profiles were also compared with the LacZ controls. The distribution of MEG3 peaks intersecting EZH2- or H3K27me3-ChIP peaks was plotted in relation to the gene region (**Fig. 3b *i***) and biotype (**Fig. 3b *ii***). Many overlapping peaks fell within the intronic regions of the genes, suggesting that MEG3 and EZH2 bind nascent transcripts and together promote PRC2 protein complex assembly [28]. In addition to the gene body, overlapping signals were observed upstream of the transcription start site (TSS) in promoter regions (**Fig. 3b *(i)***); i.e. at active genes and locations already enriched in H3K27me3 chromatin modification [23]. Top MEG3 peaks, enriched in both EZH2, and H3K27me3 ChIP signals, were identified and we obtained the maximum peak score for each gene region (**Fig. 3c**). Significant EZH2 occupancy of common targets with MEG3 hints at the possible existence of distinctive features for a group of *MEG3*-regulated genes. We identified enrichment of genes involved in the process of integrin signalling and confirmed their overlap between MEG3 and EZH2 occupancy. The *Integrin Subunit Alpha 4 (ITGA4)* showed highest occupancy by EZH2 amongst other similar genes in this signalling pathway.

We next used publicly available GEO datasets from cells depleted in MEG3, to further test the above dependencies and select mutual MEG3- and PRC2-regulated candidates. The orthologous human gene set from the microarray study **GSE73524** was obtained (from mouse C2C12 cells depleted in MEG3) as are reported in **Supplementary Table S7**. We overlapped these genes with the RNA-seq dataset **GSE71164** (from HUVECs depleted in EZH2, **Supplementary Fig. 6**), as well as gene regions from the ChIP-seq dataset **GSE114283** (reads from H3K27me3 distribution in mouse neuronal cells depleted of MEG3 *vs*. control; analyzed as in **Supplementary Fig. 5A**). Using gene orthologous analysis in <u>g:Profiler</u> we have obtained human targets from relevant mouse studies. Data intersections and overlap has identified 8 genes, one of which was ITGA4 (**Supplementary Fig. 5B**) that appeared to be dependent and regulated by MEG3 and EZH2 (**Fig. 3d, 3e**).

### EZH2 landscape in HUVECs depleted of MEG3

To next test the functional importance of MEG3-EZH2 interactions in HUVECs, we identified genomic loci with MEG3-dependent EZH2 deposition that could modulate endothelial genes expression. MEG3 knockdown has previously been linked with improved EC migration and proliferation, and we postulated this could operate through EZH2 deposition. To decipher if MEG3 depletion affected the activity of EZH2 and PRC2 complex, we employed ChIP-seq and examined the landscape of EZH2 occupancy. We immunoprecipitated EZH2, together with associated chromatin isolated from the HUVECs previously transfected with MEG3 GapmeRs (10 nM, 48h) or a scrambled control GapmeRs (Ctr) and purified bound DNA. In control cells (Ctr), EZH2 occupancy was observed at over 6000 regions, associated with total 4785 peaks. MEG3 mRNA levels were depleted by 70% using GapmeRs (**Fig. 3f *i,ii***). Depletion of MEG3 resulted in decreased levels of EZH2 at ∼10% of target regions (**Fig. 3g *i***) and was lost or reduced at a total of ∼563 loci, as represented in the Venn diagram of EZH2 targets (**Fig. 3g *ii***). The targeted promoter regions were sorted by EZH2 occupancy in Control *vs*. MEG3 deficient HUVECs, and genes enriched for common molecular functions and biological pathways were identified (**Table 3**, and **Supplementary Table S8** for list of all ChiP-seq targets in MEG3 KD HUVECs). Reduction or loss in EZH2 signal upon depletion of MEG3 was preferentially seen for genes from “biological adhesion” GO terms, or cell-cell adhesion and cell junctions; and belonging to integrin, cadherin, and wnt-signaling pathways (e.g. *ITGA4, ITGA8, ITGA11, KAT7, CDH3 etc.).* At the *ITGA4* locus, binding of EZH2-ChIP was reduced due to MEG3 depletion. Moreover, the bed intersection of EZH2 ChIP signal with MEG3-ChIRP binding sites at ITGA4 locus showed a loss of EZH2 binding due to MEG3 depletion. This finding suggests a clear dependency and mutual regulation of ITGA4 epigenetic landscape by EZH2 and MEG3 in HUVECs. Maximum peak scores of the overlapping signals at the ITGA4 promoter are given in **Supplementary Fig. 5C**.

**Table 3.** Molecular function (MF) and biological pathways (BP) of differentially expressed genes at all unique RefSeq genes within TSS ± 3 kb, sorted by EZH2 ChIP-seq occupancy in Control *vs.* MEG3-LNA GapmeR (10nM) HUVECs. Reduction or depletion in EZH2 signal obtained from ChIP-seq is seen in 10% of genes mostly belonging to biological pathways *(BP)* like biological adhesion, cell-cell adhesion, and cell junction. The total list of regulated genes is given as **Supplementary Table S8.csv file**

| Functional ChIP-gene expression analysis following MEG3 depletion |  |  |  |  |
| --- | --- | --- | --- | --- |
| ID | Source | Term ID | Term Name | padj (query_1) |
| 1 | <u>GO:MF</u> | GO:0050839 | Cell adhesion molecule binding | $8.991 \times 10^{-5}$ |
| 2 | <u>GO:MF</u> | GO:1904929 | Coreceptor activity involved in Wnt signaling pathway, planar cell polarity pathway | $1.619 \times 10^{-3}$ |
| 3 | <u>GO:MF</u> | GO:0015085 | Calcium ion transmembrane transporter activity | $2.137 \times 10^{-2}$ |
| 4 | GO:BP | GO:0007155 | Cell adhesion | $9.053 \times 10^{-5}$ |
| 5 | GO:BP | GO:0022610 | Biological adhesion | $1.083 \times 10^{-4}$ |
| 6 | GO:BP | GO:0098609 | Cell-cell adhesion | $2.892 \times 10^{-3}$ |
| 7 | GO:BP | GO:0098742 | Cell-cell adhesion via plasma-membrane adhesion molecules | $4.212 \times 10^{-3}$ |
| 8 | GO:BP | GO:0034330 | Cell junction organization | $9.316 \times 10^{-3}$ |

### MEG3 targets EZH2- and H3K27me3-regulated integrin signalling

We next identified endothelial genes that are regulated by EZH2. De-novo gene expression analysis was performed, from alignment and data mapping to human genome Hg38, to detect differentially regulated genes in EZH2-silenced HUVECs. To understand the relationship between *MEG3* and PRC2 binding we next sub-classified the ChIRP-*MEG3* targets into groups, according to EZH2 occupancy following MEG3 depletion and gene transcription activity in the EZH2*-*deficient HUVECs versus control (**Fig. 4a**). To identify targets with a differential PRC2-RNA binding, rather than those where PRC2 binds to chromatin, and we created genome-wide datasets. We identified and displayed logical relationships between genes by plotting the EZH2-FLASH RNA gene interactome (**pink**) against RNA-seq data for genes that are differentially expressed following EZH2 knockdown (**blue**, *de novo* analysed GEO RNA-seq data (GSE71164) of Scr *vs.* EZH2). This set of 1876 genes represents EZH2 deregulated RNA interactome (not bound at DNA level) that exemplify a novel unconventional function of EZH2 to bind RNA targets, specifically in line with the cytoplasmic distribution of EZH2 [29]. This also indicated to us that EZH2 could control 1876 genes-expression at the post-transcriptional level, but we have not further investigated it. Next, MEG3 ChIRP-seq peaks (**green**) are displayed against EZH2 ChIP-seq intensities over loci, following MEG3 knockdown (**yellow**). Based on their nature of interaction with PRC2 and MEG3, we sub-classified them into two types of targets, *Group 1*: MEG3-dependent EZH2 target genes at the DNA level (232 genes) and *Group 2:* at the RNA level (851 genes). In *Group1, MEG3-PRC2* DNA targets are the genes with EZH2 deposition (ChIP-seq) regulated by EZH2 (siEZH2 RNA-Seq) with low FLASH signal (no binding at the RNA level). In *Group2* are the targets with high PRC2-RNA signals (FLASH) over the genes bound to the nascent transcript, but with low occupancy of EZH2 (ChIP) and pronounced MEG3 enrichment on the chromatin. We focused on 232 targets in *Group 1*, regulated by EZH2 at the DNA level in a MEG3 dependent manner. A subset of genes had functions in cell adhesion molecule binding and again belonged to the *integrin cell surface interaction pathway.* These targets do not interact at the RNA level with EZH2 but are specifically regulated by it (**Fig. 4b** and **Supplementary Fig. 6**). Amongst the top 50 genes, ITGA4 is a potential EZH2 target, together with ITGB1. The integrin α4β1 (CD49d/CD29) is a cell adhesion receptor known to bind fibronectin (FN1, selective for the CS1 domain) and VCAM1 (Vascular Cell Adhesion Molecule-1) as a counter receptor and it exists as a dimer with ITGB1 subunit in the cell.

**Figure 4.**
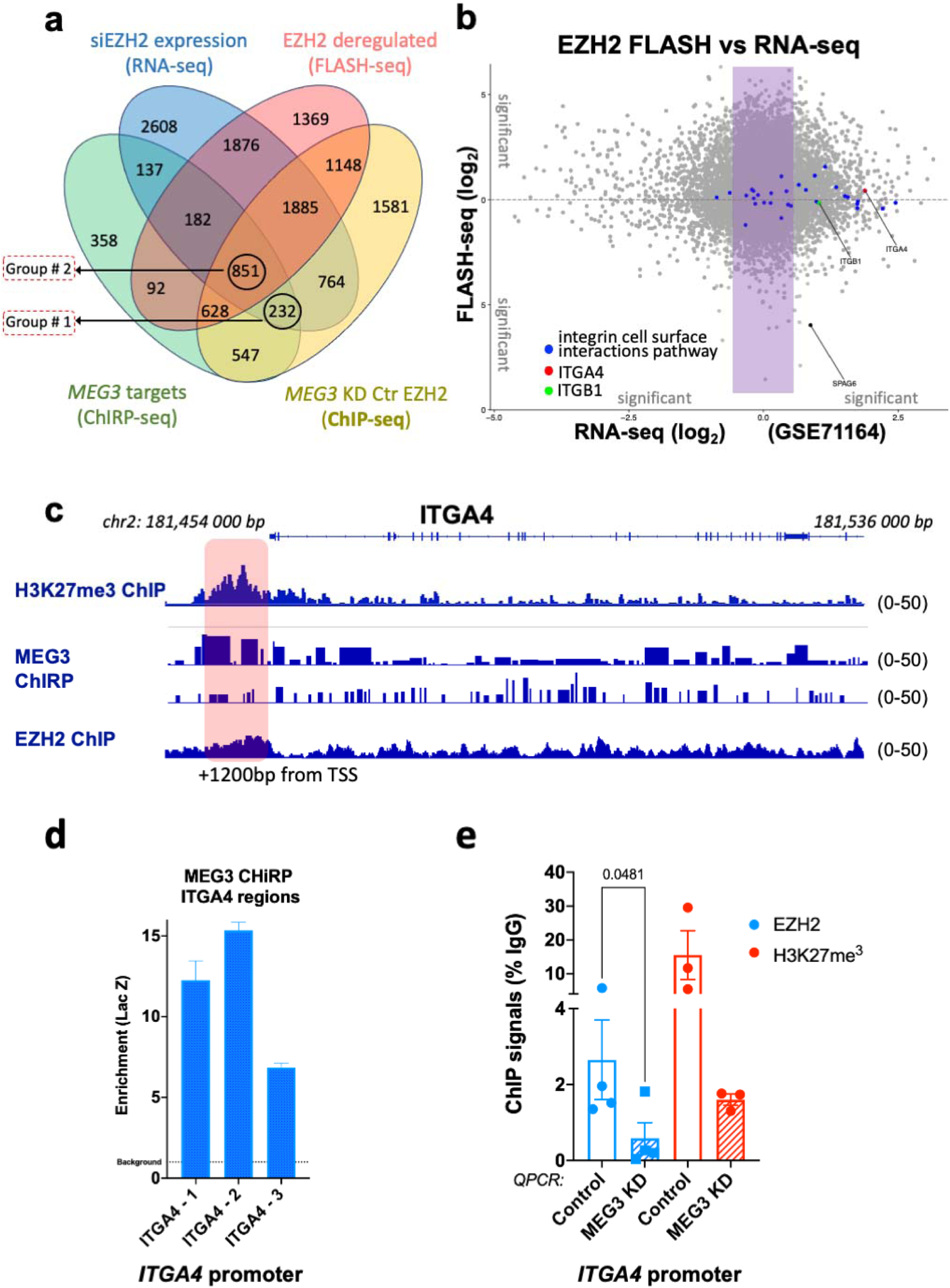
***a.*** Venn diagram showing the intersection between statistically filtered FLASH data from two biological replicates of our MEG3-ChIRP-seq-data (green), *de novo* hg38 analysed GEO RNA-seq data from siEZH2 deficient HUVECs (GSE71164, blue), and EZH2 ChIP-seq following MEG3 KD (yellow) and FLASH-seq transcriptome following EZH2 IP (pink). ***b.*** Correlation between gene expression levels and FLASH signal. Gray, expressed RefSeq genes with reproducible FLASH signal consistently detected in RNA-seq. Blue, genes with the highest RNA-seq signals and no reproducible FLASH signal belonging to integrin cell surface interaction pathway. **Red**, expressed ITGA4 gene, and **green,** ITGB1 gene, without reproducible FLASH signals. Data are from two biological replicates of each EZH2 FLASH sample and three biological replicates of EZH2 RNA-seq samples (Scr *vs.* siEZH2, GSE71164). ***c.*** Genomic tracks showing ChIRP-seq signal (MEG3 Odd, Even and LacZ) in HUVECs over ITGA4 gene only. The MEG3 binding site is located upstream of the ITGA4 gene in the promoter region, and it overlaps with the H3K27me3 signal and EZH2; as well as downstream within the ITGA4 gene body, where it overlaps with within the EZH2 signal in the intronic region of the gene. ***d.*** MEG3-ChIRP followed by qPCR, analysis of MEG3 binding region on ITGA4 in HUVECs. The crosslinked cell lysates were incubated with combined biotinylated probes against MEG3 lncRNA and the binding complexes recovered by magnetic streptavidin-conjugated beads. The qPCR was performed to detect the enrichment of specific region that associated with MEG3, peaks were related to input control and compared *vs.* the non-biotynilated control. ***e.*** ChIP-QPCR enrichment for EZH2 and H3K27me3 over ITGA4 promoter region in HUVECs depleted of MEG3 *vs.* Control.

MEG3 occupancy by ChIRP was seen within the ITGA4 promoter and in +63kb enhancer region in HUVECs. Both H3K27me3 peaks of available GEO datasets from HUVECs and *in-house* EZH2 peaks were detected in the *ITGA4* promoter region, and they all aligned with the MEG3-ChIRP peaks (**Fig. 4c**). Subsequent ChIRP-qPCR showed enrichment of MEG3 binding compared with LacZ control at three sites within the ITGA4 promoter (**Fig. 4d**). Data from one other public dataset also shows that ITGA4 is a direct target of MEG3 [30, 31]. We next performed chromatin immunoprecipitation-qPCR to quantify the levels of EZH2 and H3K27me3 repressive mark and observed enrichment of EZH2 and H3K27me3 in the promoter region of ITGA4 in control HUVECs. Assessment of enrichment following MEG3 depletion showed a reduction of both EZH2 and H3K27me3 signals in the ITGA4 promoter region (**Fig. 4e**). We conclude that the PRC2 complex both binds and is enzymatically active at the ITGA4 site. Our findings prove that the loss of MEG3 compromises the H3K27me3 landscape, as it reduced PRC2 binding to ITGA4. Accordingly, PRC2 requires binding of MEG3 for chromatin localization and methyltransferase activity in the promoter region of ITGA4.

### MEG3-PRC2 dependent regulation of ITGA4 in ECs

To gain further insight into the functional significance of the interaction of EZH2 with MEG3, we determined how pharmacological inhibition of EZH2 enzymatic activity affected cell function and gene expression. We used specific PRC2 inhibitor (A-395, an EZH2-EED inhibitor [32, 33]) of protein interaction that blocks EZH2 activity) in HUVECs and then performed chromatin immunoprecipitation-qPCR against EZH2 and H3K27me3. As expected, pharmacological inhibition of PRC2 reduced the ChIP signals of EZH2 and the associated repressive mark H3K27me3 over the ITGA4 promoter, when amplified by two sets of primers against the promoter region (**Fig. 5a**). HUVECs treated with A-395 also had increased expression of ITGA4 compared with control (**Fig. 5b**). Similarly, an increase in ITGA4 transcript level was observed following MEG3 depletion in HUVECs (**Fig. 5c**).

**Figure 5.**
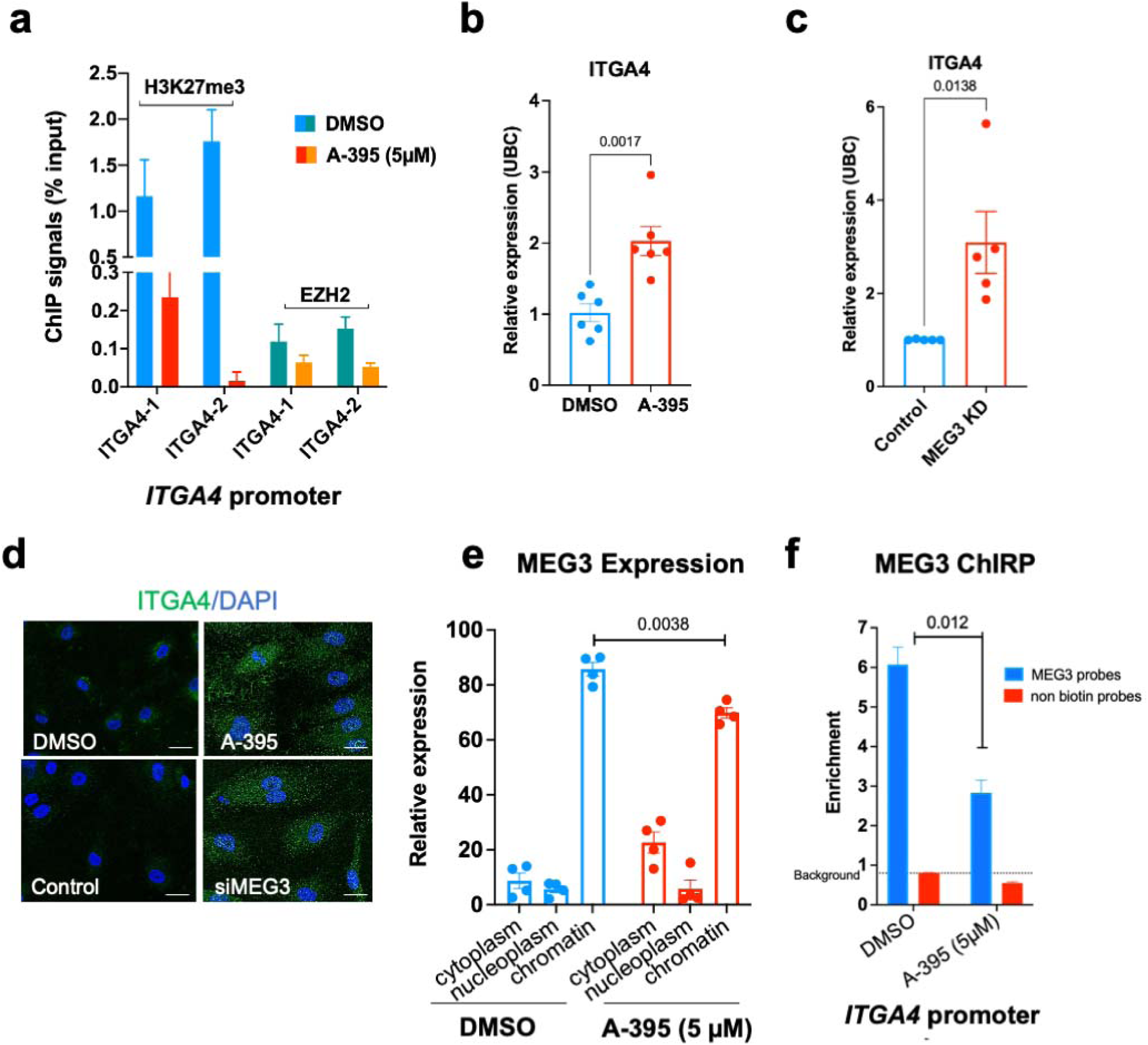
**a.** ChIP signal enrichment *vs*. 1% input for EZH2 and H3K27me3 mark over ITGA4 promoter regions in HUVECs treated with A-395 (5µM, 24h) inhibitor of PRC2 *vs.* Control (DMSO). The expression was measured using two sets of primers against the same promoter region of ITGA4. Representative graphs are average of three qPCR datasets ± SEM. **b.** ITGA4 expression in the presence of A-395 *vs*. DMSO control, *N=6* independent experiments compared using *t*-test. **c.** Measuring the expression levels of ITGA4 upon depletion of MEG3 using LNA GapmeRs (10nM, 48h), data is mean of *N=5* independent experiments (biological replicates). **d.** Representative image of immunofluorescence staining for ITGA4 protein levels in ECs treated with A-395 *vs*. DMSO, or upon MEG3 depletion like in ***b***. **e.** Intra-cellular localisation of MEG3 (chromatin associated lncRNA) between different cellular compartments in HUVECs treated with A-395 *vs.* DMSO, whereby the distribution of MEG3 has shifted upon PRC2 inhibition with A-395; from the nucleus (where it was highly chromatin bound) into the cytoplasm. Representative bars were compared by t-test and on-way Anova. **f.** MEG3-ChIRP followed by qPCR, *N*=3, analysis of MEG3 binding over ITGA4 promoter region in HUVECs treated with A-395 (5µM, 24h) *vs.* DMSO. MEG3-ChIRP HUVEC lysates treated with A-395 resulted in reduced engagement of MEG3 with ITGA4 site compared with either DMSO control or ChIRP with non-biotinylated probes. The non-biotin probes served as a negative control, and we detected the background level <1.

We used immunocytochemistry in **Fig. 5d** to identify an increase in <u>ITGA4 protein</u> (https://www.proteinatlas.org/ENSG00000115232-ITGA4) levels following MEG3 depletion or PRC2 inhibition. This further confirmed that both MEG3 and PRC2 regulate ITGA4 expression. We also assessed the subcellular distribution and found nuclear localization of chromatin associated MEG3, as previously reported. Surprisingly, we observed a shift in the cellular distribution of MEG3 when ECs were treated with A-395 inhibitor, with increased cytoplasmic occupancy of MEG3 (**Fig. 5e**). From these experiments we hypothesised that MEG3 could be removed from the chromatin. We therefore performed ChIRP in ECs treated with A-395 inhibitor of PRC2 and found that it resulted in less MEG3 enrichment at the *ITGA4* promoter region on the chromatin, compared with non-biotinylated control probes or the DMSO control (**Fig. 5f**).

### PRC2 inhibitor improves EC function via de-repression of ITGA4

Finally, we assessed EC functionality following treatment with A-395 *in vitro and in vivo.* By measuring the cell migratory capacity and determining the speed of migration, we observed that A-395 rendered ECs more capable of migrating, and increased wound closure compared with vehicle (DMSO) (**Fig. 6a**) in line with our previous reports [15]. We then addressed if the same compound could affect cell adhesion function. The interaction of cells with fibronectin (FN) is specific and mediated by integrin receptors at the cell surface [34]. The engagement of integrins by FN triggers a signalling cascade inside the cells that ultimately culminates in cell adherence and spreading. The cell index was determined as a measure of cell attachment to FN and adhesion within 3h. A-395 treatment efficiently increased the cell attachment (**Fig. 6b**). Both increased migration and adhesion capacity were reversed by *ITGA4* KD (**Fig. 6a** **and 6b**). Expression of ITGA4 was successfully reduced following *ITGA4* KD (**Supplementary Fig. 5D**). We predicted that removal of PRC2 from chromatin using A-395 would affect interactions with MEG3 *in vivo*. To test this hypothesis, we delivered subcutaneous Matrigel injection containing PRC2 inhibitor (A-395 1mg/kg) or Control to adult mice. The A-395 treatment decreased H3K27me3 (**Fig. 6c**) and increased neovascularisation (**Fig. 6d**) in the Matrigel plugs. Finally, A-395 treatment increased the percentage of vessels positive for ITGA4 compared with the DMSO (**Fig. 6e**). In summary, pharmacological inhibition of EZH2 with A-395 could increase EC function in settings like ischaemia or hypoxia, as seen in the graphical abstract (**Fig. 6f**).

**Figure 6.**
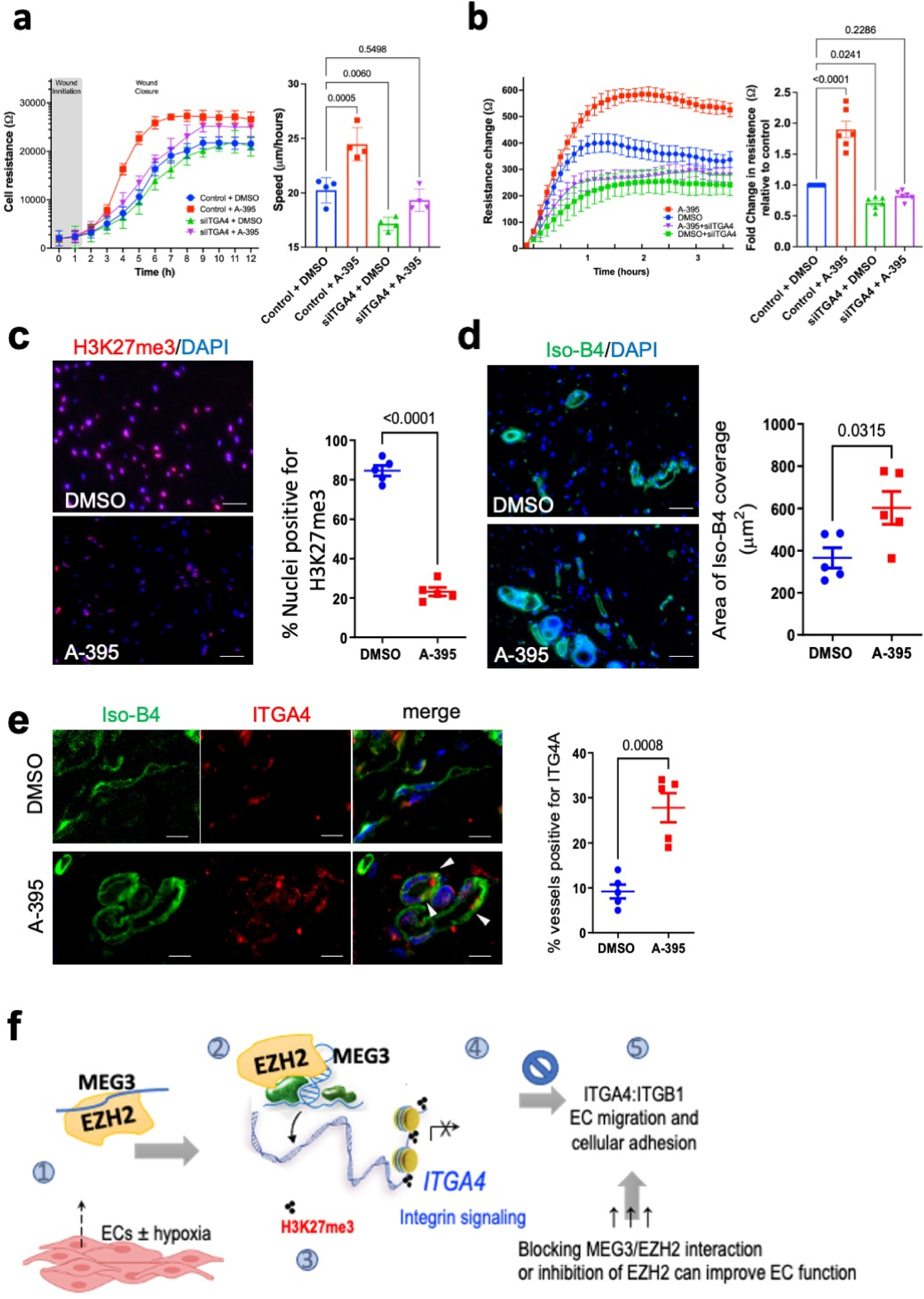
***a.*** Measure of cell migratory capacity using ECIS functional analysis in ECs treated with control or A-395 (5µM, 24h) inhibitor. Experiments were performed in duplicates (technical replicates) and four experiments were run for migration assay and six for adhesion (biological replicates). The data showing ECIS trace (left hand side) is mean ±SD as calculated by the ECIS. The graph on the right is mean±SEM with *N*=6, data was compared using ordinary one-way ANOVA with Dunnett’s multiple comparisons tests. ***b.*** Adhesion to Fibronectin, FN (20µg/ml) was used to coat the culture plates and assess adhesion of endothelial cells within 3h of ECIS assay, following cell pre-treatment with A-395, 24h. The difference in resistance change was calculated over 3h. ***c.*** Subcutaneous Matrigel plug injection (200µl) into mice (*N*=5) treated with DMSO (control, left flange) and A-395 (1mg/ml, right flange) was done for 2 weeks. Matrigel plugs were collected and processed for histology. Staining for H3K27me3 was done, displaying nuclear positivity with strong intensity in control (<0.02% DMSO in water) and the A-395 treatment decreased total H3K27me3 staining, as compared by t-test. ***d.*** Staining for arterioles was performed to assess vessel growth as angiogenesis and data was compared using Student’s t-test. The data shows increased area of staining for Isolectin B4 (Iso-B4) dye in A-395 *vs.* DMSO treated Matrigel plugs with increased neovascularization, P<0.05. ***e.*** A-395 has increased the percentage of vessels positive for ITGA4 (red) within the Isolectin B4 positive cells, compared with the DMSO using *t*-test. ***f.* Graphical abstract. *1*** Maternally Expressed Gene–MEG3 is highly expressed with hypoxia and bound to EZH2 in endothelial cells (EC) affected by ischaemic insult. ***2*** Such MEG3:EZH2 complex assembles onto the target genes to ***3*** direct the EZH2 activity to “write” H3K27me3 trimethylation repressive mark and block expression of target gene i.e. integrin alpha 4 (ITGA4) and its ability to dimerise with integrin beta 1 (ITGB1), leading to ***4*** reduced EC function as measured by adhesion and migration. Hence ***5*** targeted disruptions of MEG3:EZH2 interaction, or inhibition of EZH2 activity could increase EC function under ischaemia.

## DISCUSSION

The mechanisms of ncRNA:PRC2 interaction versus recruitment of PRC2 to pertinent chromatin targets remain an intense area of study. We combined genome-wide protein-centric and lncRNA-centric approaches to decipher the functional interaction in ECs. Here we report that the use of FLASH technique in primary ECs reliably detected both EZH2-bound RNA species (single reads), as well as RNA-RNA interactions (hybrid reads) associated with EZH2. We discovered that EZH2-binding sites in RNA contain a clearly enriched motif, indicating that sequence recognition plays a role in these interactions. Emerging evidence also suggests sequence-specific recruitment of PCR2 to chromatin, despite its apparent lack of inherent DNA binding ability [5].

Amongst top identified RNAs that directly interacted with EZH2 in ECs, we recovered lncRNA-MEG3 and nominated for functional analyses. Within MEG3, EZH2 binds stable secondary structure elements, constraining MEG3 hybrid interactions and indicating previously reported conserved pseudoknots [35]. This is consistent with *in silico* predictions referring that MEG3 is highly structured, due to high content of G – C base pairs (57%) [7, 35]. We isolated and characterised genomic binding sites for MEG3 using ChIRP and then assessed chromatin targeting by the MEG3-EZH2 complex within the EC genome. This was a pivotal task, providing biochemical evidence for direct physical and functional interactions between EZH2 and MEG3. We explicitly focused on MEG3 variants, NR_002766.2; transcription variant 1 and NR_003530, transcript variant 2, both highly expressed in ECs, to define context specific functions. The identification of secondary structures within MEG3 molecule as EZH2 binding sites suggests that MEG3 structure is the direct intermediate for PRC2 recruitment onto chromatin. Many endothelial lncRNAs mediate gene expression and guide context-specific regulation of PRC2 activity, potentially via related, structure-based interactions. It remains unclear whether other MEG3 splice variants with different MEG3 exonic organization have similar functions, exhibit different levels of interactions or MEG3 structures in HUVECs.

ChIRP with MEG3 mapped endothelial MEG3-binding loci clearly overlaps with the EZH2 occupancy in ECs. Common target genes were interrogated though gene ontology and showed enrichment for functions in angiogenesis. These genes preferentially function in endothelial cell adhesion and ECM formation; notably including integrins. Specifically, MEG3-EZH2 interaction directs PRC2 binding, serving as an epigenetic guide to deploy H3K27me3 for repression of integrin ITGA4, endothelial target. Integrins recognize a variety of proteins on the surface of other cells or in the ECM, and are therefore essential for cell adhesion, sprouting, and migration. Integrin-mediated EC adhesion to ECM is central for remodelling of cytoskeleton of ECs and sprouting [29, 36].

We discovered that direct binding of EZH2, and a concomitant association of *MEG3,* actively controls the transcription of integrin *ITGA4*. The EZH2:MEG3 interaction points coincided with *1)* the genomic region where the PRC2 is enzymatically active on ITGA4, and thus enriched in H3K27me3 and *2)* genomic regions where other PRC2 components are deposited and interact with the chromatin. A low level of H3K27me3 on ITGA4 precedes recruitment of PRC2 and deposition of further H3K27me3. Occupancy of *ITGA4* by PRC2 components is required for MEG3 binding to stimulate the enzymatic activity of PRC2. The genomic occupancy and spread of EZH2 onto *ITGA4* was reduced in the absence of MEG3. Removal of MEG3 eliminated the inhibition of EC migration and functional sprouting capacity, key processes during angiogenesis. We expose a role of EZH2 in the adhesion of endothelial cells. We also showed that expression of ITGA4 is essential for EC migration, and for regulation of stimulated cell adhesion or adhesion-dependent signal transduction. Despite the functional redundancy between different integrin subunits on endothelial cells, a specific MEG3-targeting of ITGA4 through PRC2 deposition can lead to ITGA4 transcriptional inactivation.

Both inhibition of PRC2 or MEG3 knockdown in ECs, have revealed molecular events leading to ITGA4 gene activation and improved EC function. Thus, blocking of the scaffolding role that MEG3 plays in PRC2 assembly could lead to functional benefit. Similar, context-specific, MEG3 action has previously been described in regulation of TGF-β gene in cancer cells [14]. Although we find similarities in ChIRP targets between our datasets (overlap of 2933 hits upon correlation), we did not observe the same pattern of MEG3 targeting TGF-β in ECs, implying portentous context specificity of MEG3 actions and interactors, akin to its role in myoblast identity [30]. Our results, however, suggest a novel mechanism of MEG3 ability to inactivate ECs, linking integrin-mediated EC adhesion via *ITGA4* to endure PRC2-associated chromatin changes in the nucleus; whereby epigenetic inhibition of PRC2 could derepress key genes, to enable efficient endothelial cell migration and adhesion during conditions like vascular injury and hypoxia.

Molecular mechanisms that alter the epigenetic landscape through actions of lncRNAs, or signalling (e.g. injury and hypoxia), have large implications for various pathologies entailing EC dysfunction [37]. The expression of many lncRNAs is under active transcriptional control in endothelial cells, albeit with minor regulatory roles. Thus, identification of specific function for a MEG3 with a binding capacity for chromatin-associated protein was fundamental for understanding its cell specific actions. Furthermore, generating a specific MEG3-database of associated super enhancers and other genomic regions, is a valuable resource for vascular biology that could help to unravel other MEG3-regulators and -binding partners at play in specific cardiovascular pathologies. Finally, the potential for PRC2 to exert site-specific RNA or DNA binding has opened new avenues for use of PRC2 inhibition to enhance vascular function. Future work will address how the functions of Polycomb complexes are disrupted in human diseases entailing EC dysfunction.

## METHODS

### Endothelial Cell culture and treatment and transfection

Human Umbilical Vein Endothelial Cells (HUVECs) (C-12203) were purchased from Promocell (Promocell, Germany) and cultured in the recommended growth media (EGM2, MV2) under 5% CO2 at 37 °C. Hypoxia was stimulated in Coy chamber with a gas mixture of 1% O2, 5% CO2 and 94% N2 (Coy Laboratory Products). Endothelial cell growth medium 2 (EGM2) was prepared from endothelial cell basal medium (Promocell, C-22211) and supplements (Promocell, C-39211) including Fetal Calf Serum (2%), according to the manufacturer’s instructions. Cell preparation for experiments varied in the size and number of plates used and the quantity of cells harvested. Cryopreserved HUVECs were brought into culture and incubated in complete EGM2 medium to adhere to flasks. Fresh media was added to remove residual DMSO after 24h, and cells were incubated in their first passage for 48h. Medium was replaced every 48h and cells were passaged once reaching 80-90% confluency. Cells were cultured between passage P4 and P8 for 16 duplication times (P8). All cells described in the following experiments were used at passage 5 (P5) or P6. Cells were washed with PBS, detached using trypsin/EDTA (1x, Life Technologies) and plated onto plates of desired size.

HUVECs were plated on 6-well plates at 80% confluence and left to attach overnight before pre-treating them with A-395 (Sigma, SML1923) for 24h using 5µM final concentration with 0.01% DMSO in water. For transfection experiments cells were seeded onto 12-well plate, for 24h. Next day OPTIMEM medium was added for 1h before addition of transfection complexes prepared by mixing lipofectamine iMax^TM^ (Thermofisher) with GapMers against MEG3 (10nM, Qiagen). Complexes were incubated for 6h after which EGM2 medium was added to replace complexes and optimum, and cells left for further 48h.

### RNA extraction and quantitative real time analysis

Total RNA was extracted using miRNeasy kit (Qiagen) or Monarch kit (New England BioLabs (NEB) #T2010). RNA concentration and purity were assessed using a NanoDrop2000 Spectrophotometer (ThermoFisher Scientific). For mRNA analysis RNA was reverse transcribed using LunaScript reverse transcription Kit (NEB, #E3010). cDNA amplification was performed on a QuantStudio 5 Flex (ThermoFisher Scientific) Real-Time System in 384-well plate. Luna qPCR SYBR Master Mix (NEB, M3003) was used along with specific primers to determine the expression of MEG3 and housekeeping genes using Applied Biosystems QuantStudio 5 Real-Time PCR Detection System. The comparative delta cycle threshold (Ct) value method was used for qPCR analysis. Ct values represent the number of PCR stage cycles required for cDNA of a gene of interest to be amplified enough for the fluorescent signal from the reaction to exceed a defined threshold. Full list of primers and reagents is given in ***Supplemental Table S9***.

### Subcellular Fractionation (nuclear, chromatin & cytoplasmic RNA) followed by qPCR

For subcellular isolation of RNA, HUVECs were plated onto 10cm^2^ dishes at P5. Media was changed and cell plates were treated with A-395 of DMSO for 24h. Fractionation was performed to separate nuclear, chromatin and cytoplasmic fractions as reported before [38]

#### Cytoplasmic lysis buffer

contained (0.15% (vol/vol) NP-40, 10 mM Tris-HCl (pH 7.0), 150 mM NaCl, 25 μM α-amanitin, 10 U SUPERase.IN, 1x Protease inhibitor mix, prepared freshly before use and stores on ice);

#### Nuclei wash buffer

(0.1% (vol/vol) Triton X-100, 1mM EDTA, 25 μM α-amanitin, 40 U SUPERase.IN, 1x Protease inhibitor mix, In 1x PBS) and

#### Nuclei lysis buffer

(1% (vol/vol) NP-40, 20 mM HEPES (pH 7.5), 300 mM NaCl, 1M urea, 0.2 mM EDTA, 1mM DTT, 25 μM α-amanitin, 10 U SUPERase.IN, 1x Protease inhibitor mix).

The RNA Lysis was performed with above fractions using RNA Lysis Buffer (NEB, #T2012) and following the Monarch kit (NEB, #T2010). RNA was precipitate with 60 μl of 3 M sodium acetate (pH 5.5), 2 μl of GlycoBlue (15 mg/ml) and 500 μl of RNase/DNase-free H_2_O, and extracted from each fraction, from three independent experiments as described above. Following RNA quantification using Nanodrop, reverse transcription (RT-qPCR) reaction was carried out in duplicate, with total of 300ng of purified RNA being converted to cDNA using 1X LunaScript RT SuperMix in 20 μl reactions and standard reaction conditions (25°C/2 min, 55°C/10 min, 95°C/1 min). qPCR detection was performed using the Luna Universal qPCR Master Mix (NEB, M3003) and included 1 μl of neat cDNA product as a template, with triplicate reactions for each target against MEG3, UBC or NEAT1. Results were evaluated for efficiency and lack of non-template control (NTC) amplification.

To study the cellular localization of MEG3, the qPCR levels MEG3, NEAT1 and UBC in the nucleus and cytoplasm were plotted as percentage, defined as: *percentage nucleus* = (MEG3 quantified in RNA nuclear fractions)/(NEAT1 quantified in total fraction RNA) and *percentage cytoplasm* = (MEG3 quantified in RNA cytoplasmic fractions)/(UBC quantified in total fraction RNA), using NEAT1, UBC and 18S as loading controls of nucleus, cytoplasm, and total cellular fractions, respectively.

### UV crosslinking of cells

For plates to be treated with UV, 15cm culture dish with confluent HUVECs were washed in ice-cold PBS, PBS was removed before placing cells on a bed of ice and crosslinking with 400 mJoules/cm^2^ UVB frequency λ = 254 for 2 min in a CL-1000 UltraViolet Crosslinker UVP Stratalinker® (Stratagene). Cells were washed twice in ice cold PBS (300RCF, 5 minutes, 20°C) and scraped to collect the pellet (16,000RCF, 5 minutes, 4°C). During the final wash, cells were resuspended in 1ml PBS and transferred into 1.5ml Eppendorf then centrifuged again and pellet was immediately used in fractionation step or cell lysis.

### <u>F</u>ormaldehyde/UV assisted cross-linking <u>l</u>igation <u>a</u>nd <u>s</u>equencing of <u>h</u>ybrids (FLASH-seq)

The approach used is similar to that reported for CLASH [39], whereas RNA-protein interactions were captured in growing cells using UV crosslinking and antibodies for affinity purification of endogenous RNA-protein complexes instead of FLAG-tagged protein [21]. Brief formaldehyde crosslinking is used during purification step, to stabilize binding of the covalent bait protein-RNA complex to the protein A beads. This step allows column washes under highly denaturing conditions, hence assuring reliable downstream hits.

HUVECs were cultured as described, grown to 80% confluency and UV crosslinked as above to obtain the cell pellet. Pellet was lysed in ice-cold TM150 buffer (20 mM Tris-HCl pH 7.4, 150 mM NaCl, 0.4% NP-40, 2mM MgCl2, 1 mM DTT, protease inhibitors (Roche, complete, EDTA-free), RNAse Inhibitor (Promega)). 10µl of RQ1DNAse (Promega) were added, the samples were mixed by pipetting and incubated for 10 min at room temperature to break genomic DNA. Lysates were centrifuged in Eppendorf mini centrifuge (14000rpm, 4°C, 10 min) and supernatant was collected.

#### Affinity purification

Cell lysates were incubated with 20 µl of anti EZH2 antibodies (D2C9) or rabbit IgG (see **Supplementary Table S9**.) for 2h at 4°C, then 50 µl Protein A beads (LSKMAGA02, Millipore) were added for another 60 min. The ribonuleoprotein (RNP) complexes bound to the beads were washed twice with PBS-WB buffer (PBS, +150mM NaCl, 2 mM MgCl2, 0.4% NP-40), and were treated with 0.5U RNaseA+T1 mix (RNace-IT, Stratagene) for 10 min at 20°C, then washed again twice with PBS-WB buffer and once in PBS. These complexes were cross linked on beads in 0.1% formaldehyde (PFA, Life Technologies #28908) in PBS for 2 min, then PFA was quenched by addition of glycine to 0.2M and Tris-HCl pH=8 to 0.1M. Crosslinked complexes were subjected to 3x denaturing washes in UB (50 mM Tris pH=7.4, 2M UREA, 0.3M NaCl, 0.4% NP-40) and 4x washes with PNK buffer (50mM Tris-HCl pH 7.5, 10mM MgCl2, 50mM NaCl, 0.5% NP-40, 1mM DTT) to remove non-specific interactions. Subsequent steps were identical for FLASH as described for CLASH [39].

#### RNA end modification and linker ligation

The complexes were treated with TSAP phosphatase (Promega) for 40min at room temperature. Between all enzymatic reactions immobilized complexes were washed once with UB and three times with PNK buffer. Phosphorylation of RNA was carried out with 40 units T4 PNK (New England Biolabs), first with P32 labelled ATP for 45 min, then 20 more min with 1 mM cold ATP at room temperature. The reactions should provide 5’ phosphates needed for downstream ligations.

Protein-bound RNA molecules were ligated together and with 3’ linker (1 μM miRCat-33, IDT), overnight using 40 units of T4 RNA ligase 1 (New England Biolabs) at 16°C. This reaction created RNA hybrids and single RNA molecules ligated to miRCat linker.

Then barcoded 5′ linkers (final conc. 5 μM; IDT, one for each sample) were ligated with RNA ligase 1 in its buffer with 1mM ATP for 3-6h at 20°C; see **Supplementary Table S9**. The complexes were eluted off the beads by boiling for 3 min in NuPAGE protein sample buffer with 100 mM Tris-HCl, 1%SDS, 100 mM ME (β-mercaptoethanol).

#### SDS-PAGE and Transfer, Proteinase K Treatment, RNA Isolation and cDNA Library preparation

Eluted RNA-protein complexes were resolved on a 4%–12% Bis-Tris NuPAGE gel (Life Technologies then transferred to nitrocellulose membrane (GE Healthcare, Amersham Hybond ECL). The membrane was exposed on film (Amersham) at -70C. The radioactive bands corresponding to the EZH2-RNA complexes were cut out, and then incubated with 150μg of Proteinase K (0.1µg/ml final, Qiagen #19131) for 2h at 55°C. The RNA was extracted with phenol-chloroform-isoamyl alcohol (PCI) mixture and ethanol precipitated overnight. The isolated RNA was reverse transcribed using miRCat-33 primer (IDT) with Superscript III Reverse Transcriptase (Life Technologies) and RNA was then degraded by addition of RNase H (New England Biolabs) for 30 min. cDNA was amplified using primers P5 and primer PE_miRCat_PCR and TaKaRa LA Taq polymerase (Takara Bio). Reverse Transcription (RT) was carried out with RT primer miRCat-33 primer (IDT) CCTTGGCACCCGAGAATT. Then we carried out 21 cycles of PCR (PE_miRCat_PCR) with the following PCR primers:

P5 AATGATACGGCGACCACCGAGATCTACACTCTTTCCCTACACGACGCTCTTCCGATCT
PE_mircat_reverse CAAGCAGAAGACGGCATACGAGATCGGTCTCGGCATTCCTGGCCTTGGCACCCGAGAATTCC

The product was gel purified, and samples pooled before submitting for sequencing. In total 20 µl of 3 ng/ µl mixed sample was sent for sequencing with ∼0.3 ng/µl of each individual sample. The final library was of the following form:

5’- AATGATACGGCGACCACCGAGATCTACACTCTTTCCCTACACGACGCTCTTCCGATCT===INSERT===T GGAATTCTCGGGTGCCAAGGCCAGGAATGCCGAGACCGATCTCGTATGCCGTCTTCTGCTTG-3’

Barcoded cDNA libraries from each sample were pulled together and subject to high-throughput Illumina sequencing (PE100), with Bejing Genomics Institute (BGI).

#### FLASH sequencing data processing

Duplicate reads caused by PCR amplification were removed using the pyFastqDuplicateRemover.py tool from the pyCRAC suite [40]. Strand specific EZH2 reads were assigned to all detected RNA biotypes and distribution analysis of single hits was performed to obtain genomic location. Trimmed and filtered FLASH single hits were aligned to the human genome (hg38) using Novoalign 2.07 [41], and overlaps between aligned reads and genomic features were calculated using the pyReadCounters.py tool from the pyCRAC suite. Differential binding in FLASH data between EZH2 and IgG control experiments, each of which were represented by two biological replicates, was calculated using DESeq2 [42]. The RNA-RNA hybrids were identified using the hyb pipeline [43], and MEG3 hybrid fragment sequences were extracted using hybtools [44]. The outputs of hyb pipeline contained a file with coordinates of all the chimeras (“.hyb”) and a file with the folding analysis of chimeras (“.viennad”), and all other output file formats are described in the hyb pipeline documentation. Enriched motifs in hybrid sequences were identified using MEME [45]. Enriched motifs in the sequences of FLASH single hit clusters were identified using MEME with motif logos and *E*-values (statistical significance from MEME) for top significant k-mers (4-8nt in length) [45]. Differentially expressed lncRNAs were obtained from FLASH-seq by mapping against <u>Ensembl</u> gene annotations. We associated lncRNAs to neighbouring genes. Overlapping reads were merged into clusters using the pyClusterReads.py tool from the pyCRAC suite [40], and the sequences of these clusters were obtained using bedtools getfasta [46].

### UV-RNA immunoprecipitation (RNA IP)

HUVECs (2×10^6^ cells perp late) were harvested from 2 15cm-tissue culture plates following UV crosslinking as described above. Crosslinked pellet was lysed in ice cold RIPA Lysis buffer pH 8.0 (1mL/pellet) (50mM Tris pH 8,0, 150mM KCl, 0.1% SDS, 1% Triton-X, 5mM EDTA, 0.5% sodium deoxycholate) with addition of the following (per 1mL): 100U/ml RNAseOUT, 0.5mM DTT and 1X PIC as per suppliers’ instructions and rotated for 30 min at 4 °C. Resuspended lysates were sonicated as described above and supernatant collected following spin in microcentrifuge (16000rpm, 20 min, 4 °C). To pre-clear the lysates 25 µl of Protein G Dyneabeads (10003D, Life Technologies) were added per IP condition for 30min at 4°C, then removed. At this stage we removed a 10 µL aliquot for RNA INPUT. Immunoprecipitation with pre-cleared UV-crosslinked lysates of ECs has proceeded overnight at 4°C using antibodies against repressive chromatin (EZH2 and H3K27me3, see ***Supplemental Table S9***), IgG or mock control (no antibody), as described for ChIP. Beads were captured on a DynaMag magnet (Thermofisher, 12321D) and washed in 500 µl ice cold Native Lysis buffer (150mM KCl, 25mM Tris pH 7.5, 5mM EDTA, 0.5% NP-40 with fresh addition of 0.5mM DTT, 1X PIC, 100U/mL RNaseOUT) to wash off the unbound material. Three RIP washes were performed at 4°C and beads pelleted (2,500 rpm, 30 s), followed by a final wash of 500 µl with Tris EDTA (10 mM Tris.HCl pH 7.5/1 mM EDTA). The beads or input samples were then resuspended in TRIzol (1ml) vortexed vigorously for 10 sec and incubate at RT for 10 min. Samples were usually store at -80°C before further RNA extraction using Monarch kit (NEB, #T2010) following manufacturer’s instructions. RNA was eluted with nuclease-free water (e.g. 20 μl) quantified and used in qPCR analysis that was performed in triplicate.

### Sonication of chromatin

UV crosslinked lysates from FLASH and RIP were sonicated (6X 2mins ON/30secs OFF) to fragment DNA to 200-700bps, and centrifuged (15mins) using Bioruptor sonicator (Diagenode). Glutaraldehyde crosslinked (1%) lysates from ChIRP were sonicated at Covaris E220 Evolution Focused-Ultrasonicator System using a water at temp range 5-7 °C and following protocol: peak power 140 W, duty factor 10%, 200 Cycles per burst for 60 sec. Total of 30-35 cycles were used for cell lysates prepared from 150mg of crosslinked HUVEC pellet. Samples were then centrifuged (16,000RCF, 15 mins at 4°C) and 10μl was supernatant was removed from each condition for DNA isolation following de-crosslinking. DNA was eluted by adding 150μl elution buffer and shaking (37°C, 30mins, 900rpm) and 15μl proteinase K (10mg/ml, Sigma, P4850) was added per sample and shaking repeated (50°C, 45mins, 900rpm). DNA was purified using QIAquick® PCR Purification Kit. Fragment size was determined by resolving samples on 1.5% agarose gel with 1kb and 100bp ladders for reference. The gel was imaged, and fragments visualised using UVidoc HD6 system (UVitec).

**Chromatin isolation by RNA purification (ChIRP)** was performed as described previously [47] using two sets of antisense oligonucleotide biotinylated DNA probes targeting MEG3 and lacZ, that were designed at http://www.singlemoleculefish.com/designer.html, some of which already published and validated. 3’-biotinylated antisense MEG3 DNA oligonucleotide probes were incubated with chromatin lysates from HUVECs. Biotinylated LacZ probes served as negative controls. Before, cells were fixed with 1% glutaraldehyde (Sigma-Aldrich) and the cross-linking reaction was quenched with glycine (0.125mM). The crosslinked cells were pelleted (>100mg) and stored at -80 °C until use. Next, 100-150mg pellet was pulverized in liquid nitrogen and chromatin prepared as reported before [48]. 1% lysate/sample condition was saved as input. 10%/90% sample were used for RNA/DNA extraction respectively. Chromatin complexes were purified using RNA pulldown with streptavidin-labelled C-1 magnetic beads (65001, Invitrogen). Subsequent stringent washes of beads:biotin-probes:RNA:chromatin were performed as described already [49]. MEG3-bound DNA and proteins were eluted off the beads using nuclear hybridization buffer (0.75 M NaCl, 50mM Tris.HCl pH 7.0, 1mM EDTA, 1% SDS, 15% Formamide and Water, with 1mM AEBSF, 0.01x vol protease inhibitor cocktail (Sigma P8340) and 0.01x vol RNase Inhibitor (Sigma R1158) added fresh just before use) and incubated for 60 min at 37°C. After the incubation, beads were collected and washed four times with 500 ml of ChIRP wash buffer (2X SSC, 0.5% SDS, add fresh 1mM AEBSF and 0.01x vol RNA inhibitor). After the last wash, beads were deposed in elution buffer (50 mM NaHCO_3_, 1% SDS, 200 mM NaCl) with 10ul of Proteinase K (10mg/ml, Sigma P4850) and reversal of crosslinking was performed at 65 °C for 3h. DNA was eluted with a cocktail of 100 mg/ml RNase A (Sigma-Aldrich) and 0.1 U/ml RNase H (Ambion) and recovered using Monarch kit and used for library construction. RNA was TRIzol-extracted and purified. The quality and RNA was assessed using Agilent RNA 6000 Pico Kit and Agilent Bioanalyser, obtaining the quantification information. Qubit Hight Sensitivity DNA Kit (Q32851, Life Techologies) was used to obtain total DNA concentration and DNA peak distribution and fragments or DNA 1000 allowing a correct sizing and quantification analysis. The input and odd and even probes samples were sequenced individually.

#### ChIRP-seq data processing

Initial processing and alignment of sequenced ChIRP-Seq data was performed by BGI, who performed initial fastq filtering using SOAPnuke [50], aligned the filtered reads to the human genome (hg19) using SOAP2 [51], and called peaks using MACS 1.4.2 [52]. ChIRP-Seq peaks for duplicate probes obtained from BGI in bed format were lifted over from hg19 to hg38 using liftOver [53], and concatenated. Genomic coverage of ChIRP-Seq peaks was calculated using bedtools merge [46] and bedGraphToBigWig [53]. Profiles and heatmaps around the transcription start site (TSS) of mRNAs were created using the ComputeMatrix, plotHeatmap, and plotProfile tools from the deepTools suite [54].

### Chromatin immunoprecipitation followed by sequencing or qPCR after MEG3-KD

ChIP-qPCR experiments were performed as previously described [15] with modifications. A total of 2 × 10^6^ HUVECs were seeded for each condition and transfected with 10 nM scrambled LNA (locked nucleic acids) GapmeR control (Cat. No. 339515) or phosphorothioate antisense standard GapmeRs MEG3-lncRNA for 48h (Cat No. 339511, Qiagen). HUVECs were then harvested and fixed with 1% formaldehyde solution (methanol free, Thermo Fisher Scientific) at a volume of 0.5 mL for every 1 × 10^6^ cell for 10 min at room temperature with slow rotation. Following sonication as described, samples were immunoprecipitated using EZH2 (D2C9) XP(R) Rabbit mAb, (5246S Cell signalling technology), Tri-Methyl-Histone H3 (H3K27me3) (C36B11) Rabbit mAb (9733S, CST) antibodies or IgG control (Normal Rabbit IgG, 2729S, CST) and captured on beads using Protein G Dyneabeads (10003D, Life Technologies). On beads crosslinked chromatin complexes were reversed, and DNA purified using QIAquick® PCR Purification Kit. The comparative delta cycle threshold (Ct) value method was used for qPCR analysis with positive and negative primer controls to validate the binding.

#### Overlap between ChIRP targets and EZH2-binding sites, and bioinformatically predicted MEG3-regulated enhancers

**ChIP-Seq** extracted bed files of histone mark H3K27me3 were downloaded from GEO <u>GSM733688</u> and <u>GSM945180</u>. The bed files of public ChIP-seq database for epigenetic regulator EZH2 in HUVECs was downloaded under accession number <u>GSE109626</u>. All intersections with called peaks from ChIP-Seq data obtained from GEO were calculated using bedtools intersect [46]. All intersections of extracted peaks in common from ChiP-seq and ChIRP-seq were calculated using bedtools intersect [46]. Maximum peak scores for ChiRP-Seq peaks that overlap genomic features were calculated using bedmap from the bedops suite [55].

To map the enhancers within the MEG3 peaks from ChIRP-seq, we used the genomic regions of HUVECs and compared them with the repository of predicted enhancers from <u>DENdb</u>: database [56]. Enhancer co-ordinates were available from http://www.cbrc.kaust.edu.sa/dendb/ (10 Jun 2020) and converted to a bed file.

MEG3 reads overlapping the enhancer regions we assessed using bedtools. Detailed processing of data is outlined in **Supplementary Fig. 4**. We attributed MEG3-enhancer sites to the nearest genes via their functional pathways and for top identified genes the gene ontologies are given in **Table 2**.

### *De Novo* Analysis of RNA-seq data

Raw SRA files of RNA-seq data were obtained from GEO <u>GSE71164</u>, converted to FASTQ files and aligned using TopHat2 (v. 2.0.3) [57]. De novo analyses was performed and data aligned to human genome build Hg38 via STAR [58] and DESeq2 [42], using a human genome database from Ensembl release 77 (www.ensembl.org). GTF files were generated as described before [59]. Statistical analysis of the differential expression of genes was performed using edgeR. Genes with False Discovery Rate (FDR) for differential expression lower than 0.01 were considered significant.

### Pharmacological inhibition of PRC2

The solution of A-395 inhibitor (Sigma, SML1923) was prepared as per manufacturer’s instructions. A-395 binds to the H3K27me3 binding pocket of EED to allosterically inhibit PRC2. For *in vitro* assays it was used at 5µM final concentration with 0.01% DMSO in water from the 50µM stock, whereas for *in vivo* studies with Matrigel the final concentration was 1 mg/ml, used from a stock of 100 mg/ml with 0.02% DMSO in water. Aliquoted stock solutions were kept at +4°C until needed for experiment.

### *In vitro* functional studies

HUVECs were pre-treated with A-395 (5µM) in 12-well plates, for 24h or transfected with siRNA for ITGA4 (20nM, Thermofisher). Endothelial cells were plated in fibronectin-coated (20µg/ml)8W10E+ electrode chamber array. Cell adhesion was continuously recorded for further 7h using ECIS Z-Theta system (Applied Biophysics) with associated software as reported before [60] and then resistance changes were calculated. Migration was analyzed using ECIS chip array (8W1E) coated in fibronectin-gelatin coating mix (10µg/ml fibronectin/0.01% gelatin). The migration speed was calculated in micrometers per hour.

### *In Vivo* Matrigel Plug Assay

Experiments involving mice were covered by the project and personal licenses issued by the UK Home Office, and they were performed in accordance with the Guide for the Care and Use of Laboratory Animals (the Institute of Laboratory Animal Resources, 1996) and in accordance with Animal Research Report of In vivo Experiments (ARRIVE) guidelines. C57BL/6N (B6N) mice (male, 10 weeks old) were subcutaneously injected into the groin regions with 300μL Matrigel (Corning) containing recombinant mouse basic FGF (PeproTech, 250 ng/mL) and heparin (Sigma, 50 U/mL) mixed with vehicle (DMSO) or A-395 (5µM). After 21 days, mice were sacrificed, and the Matrigel plugs were removed and fixed in 4% paraformaldehyde and processed for histology. Paraffin cross-sections were blocked with normal goat serum, incubated with anti-H3K27me3 (CST) and anti-ITGA4 (1:200, Peprotech) primary antibody or biotinylated Isolectin-B4 (ThermoFisher) overnight at 4°C, and then incubated with AlexaFluor 555-conjugated anti-rat IgG antibody or streptavidin AlexaFluor 488 conjugated (1:1000) (ThermoFisher). High power fields were captured (at 400X) and the number of vessels per field were counted. At least 30 randomly chosen fields were evaluated per sample, in a blinded experiment. IsolectinB4-positive vessel area was quantified using ImageJ software and expressed per square micrometres.

Subcutaneous Matrigel plugs (200µl) were injected into mice bilaterally beneath the flange and kept for 2 weeks. Matrigel solution was prepared on ice and dissolved with water and DMSO (control, left flange) or A-395 (1mg/ml, right flange) and injected one at the time with maximum 300µl volume in each mouse. After 2 weeks Matrigel plugs were collected and processed for histology. Staining for arterioles was performed to assess vessel growth as angiogenesis.

### Statistical analysis

Unless otherwise stated each experiment was performed three times (biological replicates). All statistical comparisons between two groups or multiple groups were performed using GraphPad Prism 9 software, respectively. *P* values were also calculated after performing two-way analysis using Student *t*-test or ANOVA test followed by Bonferroni post hoc test or Dunnett’s correction as indicated in Figure legends. Statistical significance was determined when *P* ≤ 0.05, and *P*>0.05 was not significant. If not alternatively specified, all error bars represent mean values with SEM.

## AVAILABILITY

All sequence data from this study, Genome maps, Bedgraph and Gene Transfer Format (GTF) files generated from the analysis of the Hfq CLASH, ChIRP-seq, ChIP-seq and RNA-seq data are available from University of Edinburgh Data repository at https://bifx-core3.bio.ed.ac.uk/hyweldd/tmitic and are currently being made available through https://zenodo.org/. All other relevant data supporting the key findings of this study are available within the article supplementary files and all computational analysis processing pipelines are available from https://github.com/hyweldd. The hyb pipeline for identifying chimeric reads and the hybtools package for downstream identification of first read in a pair is available at https://github.com/hyweldd/hybtools.

## ACCESSION NUMBERS

**ChIP-Seq** extracted bed files of histone mark H3K27me3 for chromatin state in motor neurons were downloaded from <u>GSE114283</u>. Peaks unique to Ctrl *vs.* MEG3 KD were assigned to positions associated with promoters and gene lists that were obtained from Ensembl **Biomart**, geneset *version*97 with added 2000bp upstream [61]. Gene region list was submitted to **GREAT** (Genomic Regions Enrichment of Annotations Tool) to compose associated mouse gene list [62]. **DAVID** enrichment tool and clustering were performed [63] and computational analysis pipeline is described as outlined in **Supplementary Fig. 5A. RNA-seq analysis** employed RNA sequences extracted from the GEO dataset **<u>GSE71164</u>**. **Microarray** dataset from identification of differentially expressed genes in murine C2C12 cell lines after MEG3 knockdown was extracted from <u>GSE73524</u> dataset. Using <u>GEO2R</u> two or more groups were assigned and we compared (Ctrl *vs*. MEG3 KD) to obtain top differentially expressed genes as instructed https://www.ncbi.nlm.nih.gov/geo/info/geo2r.html#how_to_use [64]. List of identified targets and human orthologues genes are given in **Supplementary Table S7**.

### Data processing and software analysis

The gene-type distribution of reads was revealed using *Pavis* online software, PAVIS [65] and gene ID overlap done using JVenn [66]. GO pathway analysis was performed on determined protein-coding genes, using Panther pathways [67], g:Profiler (ver 0.6.7.) [68] and Enrichr [69, 70]. The g:Profiler database Ensembl 103, Ensembl Genomes 50 (build date 2021-04-14) were used. Only GO terms with *P* value < 0.05 were used for further analysis. List of mouse protein-coding genes was converted to human orthologous gene ID that were obtained using g:Orth https://biit.cs.ut.ee/gprofiler/orth or converted using https://biit.cs.ut.ee/gprofiler/convert, and used in downstream representation. The python pyCRAC v1.5.2. [40], software package was used for analysing the data, available from ECDF gitlab repository https://git.ecdf.ed.ac.uk.

## SUPPLEMENTARY DATA

Supplementary Data are available online.

## ACKNOWLEDGEMENTS

T.M. is the senior author of this work and is responsible for the conception, study design, resources, experimental investigation, data collection and analysis. T.M. wrote the manuscript and revised drafts.

T.D. performed FLASH experiments and prepared library for sequencing. A.N performed ChIRP, molecular biology, *in vitro* and ex-vivo experiments and analyses. E.P. and A.C performed fluorescent staining and provided essential equipment and infrastructure for imagining analyses. A.T. performed *in vivo* experiments and analysed the data. P.M. contributed to analysis of *in vivo* data. H.D.D. performed bioinformatics analyses, P.G., A.M. and J.R. contributed to bioinformatic analysis; T.M., A.C., A.H.B., D.T. and P.M. advised on experimental design and provided essential equipment and infrastructure. D.T., A.H.B. contributed with resources, advised on experimental design, and contributed to manuscript editing and discussions.

## LIST OF SUPPLEMENTARY DOCUMENTS

**Supplementary Table S1**. **FLASH-seq hits**. List of all EZH2-FLASH targets from two replicates, with biotypes, raw reads, normalized DeSeq2 and log_2_FC between EZH2 and IgG precipitated RNA **.csv file**

**Supplementary Table S2. MEG3 FLASH single hits.** List of MEG3-RNA chimeras identified in both fragments of EZH2-FLASH replicates, single hits showing unique MEG3:RNA base-pairings that interact with EZH2 (MEG3 single hits fasta) **.csv file**

**Supplementary Table S3. MEG3 FLASH hybrids.** List of accompanying MEG3:MEG3 base-pairing (hybrids) with *nucleotide* position of the reads within MEG3 gene corresponding to exon 3-7. Hybrids represent mostly likely base pairings form with the folding predictions as per binding energy dG .*Viennad* file

**Supplementary Table S4**. **MEG3-ChIRP targets.** List of all combined MEG3 targets **.csv file**

**Supplementary Table S5**. Summary of MEG3-associated HUVECs enhancers and super enhancers with nearby genes .**csv file**

**Supplementary Table S6**. MEG3-ChIRP overlapping with published GEO ChIP of H3K27me3 and EZH2 with signal maximum peak height **.csv file**

**Supplementary Table S7**. Summary with list of all orthologues human genes to mouse C2C12 data. **csv file**

**Supplementary Table S8. ChIP-seq regions sorted by EZH2 occupancy in Control *vs*. MEG3 deficient HUVECs .csv file**

**Supplementary Table S9.**
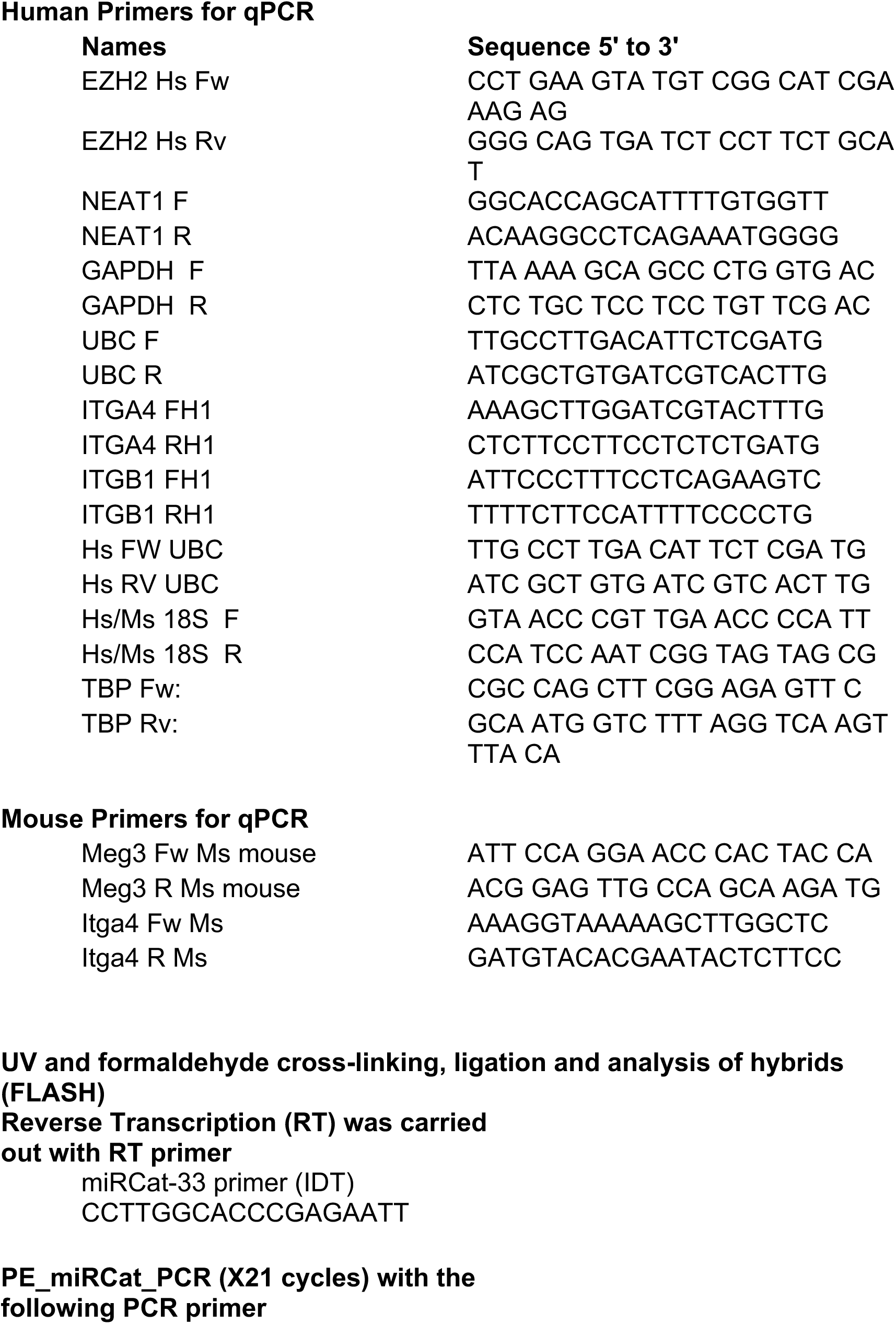
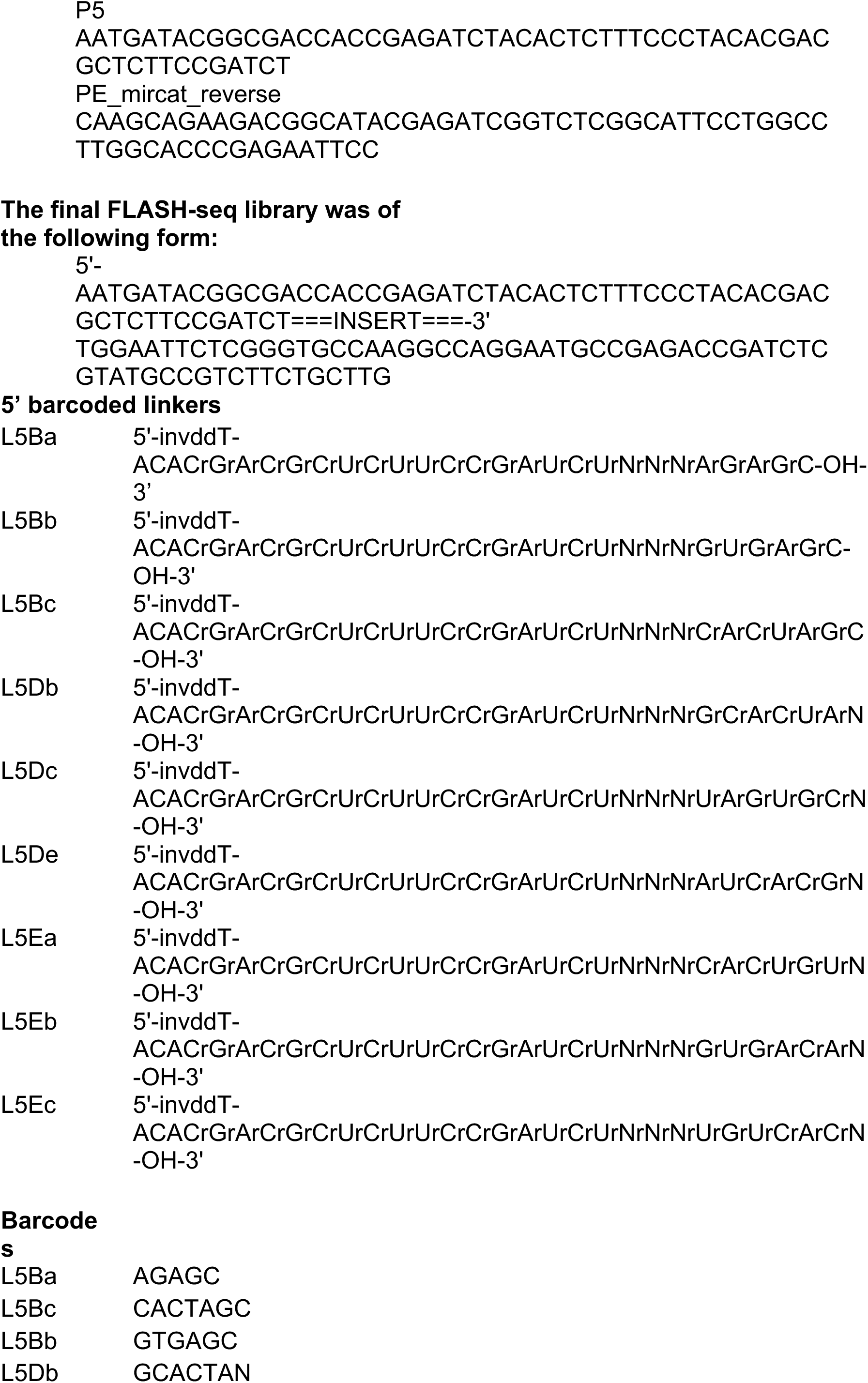
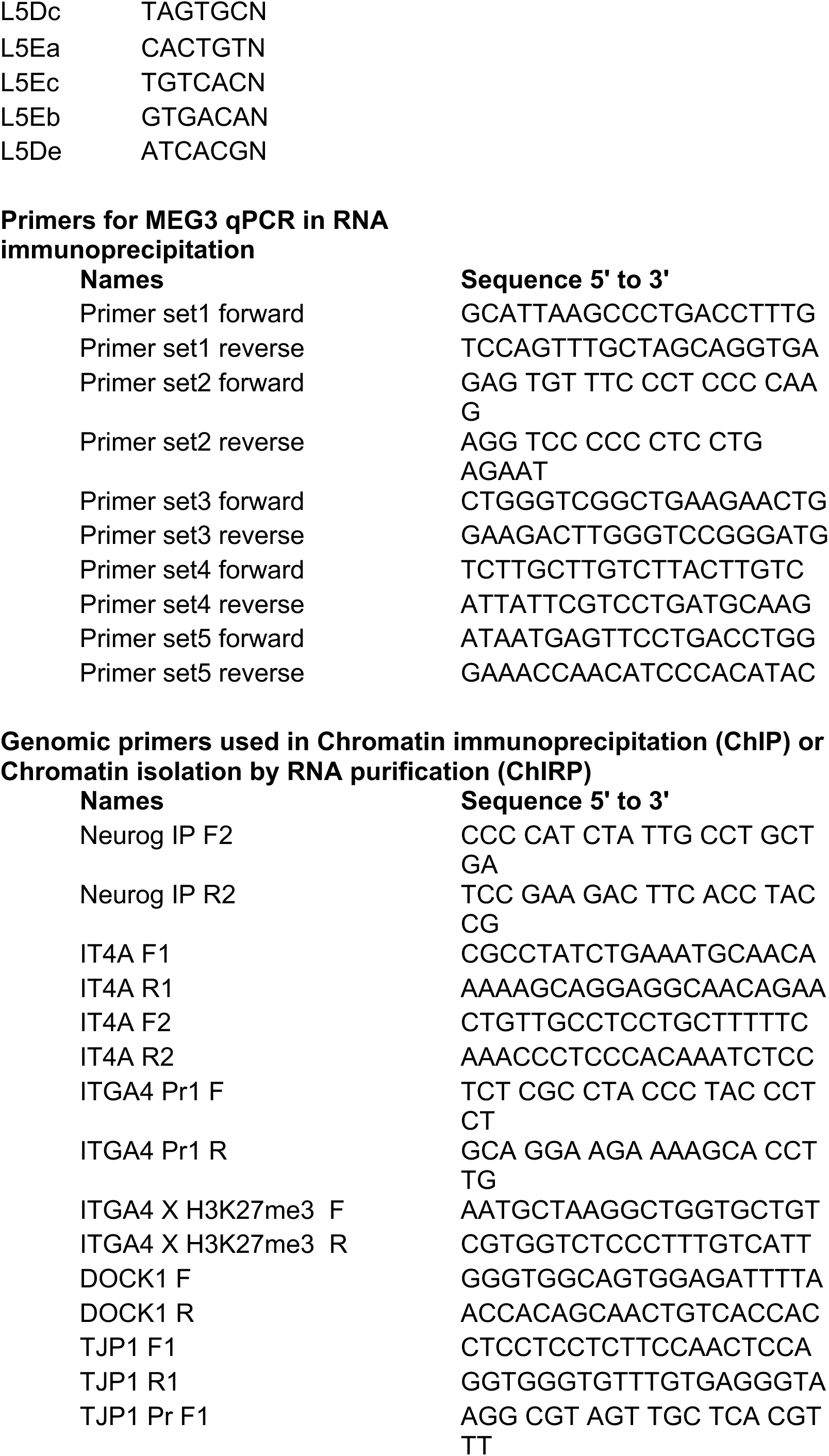
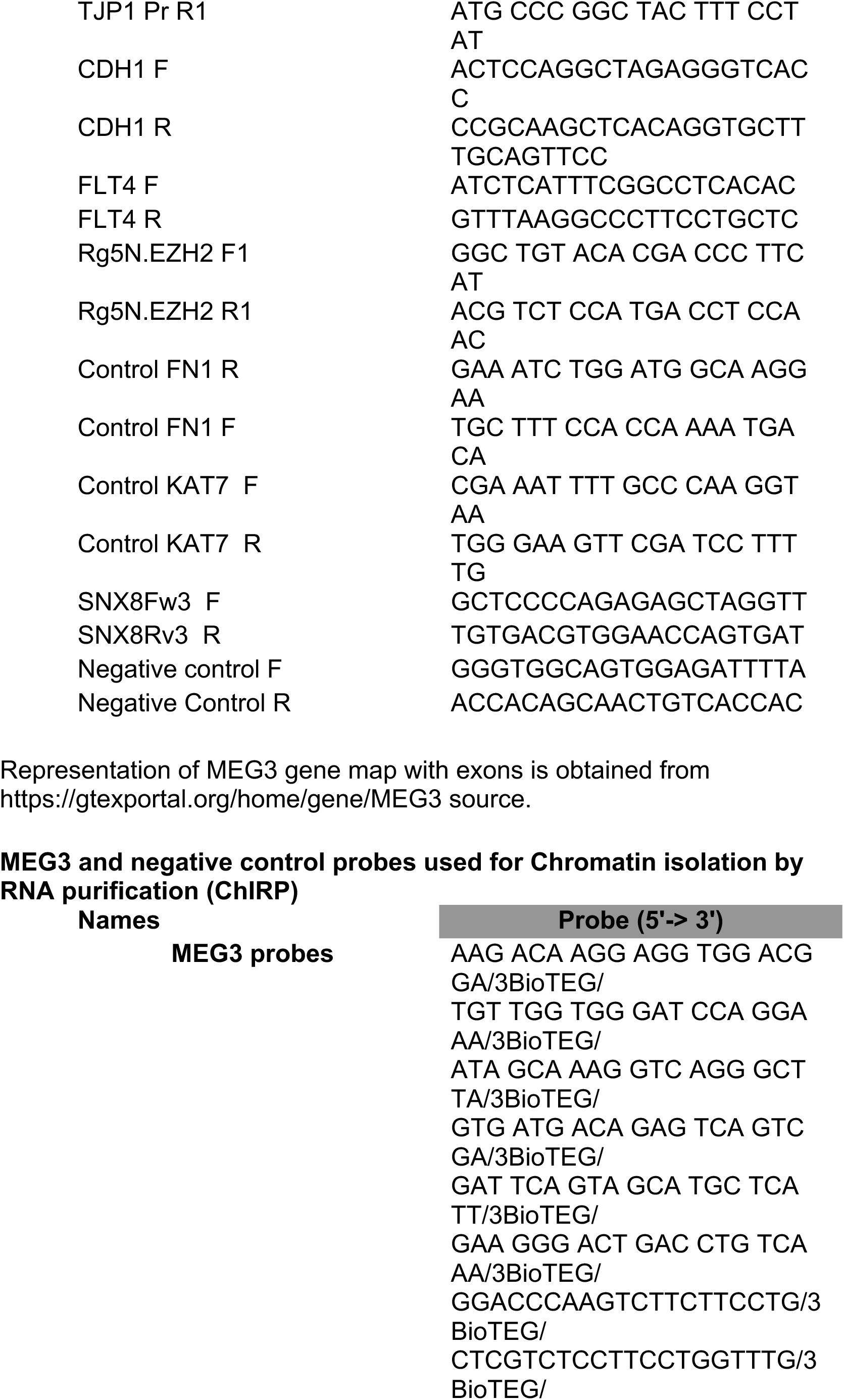
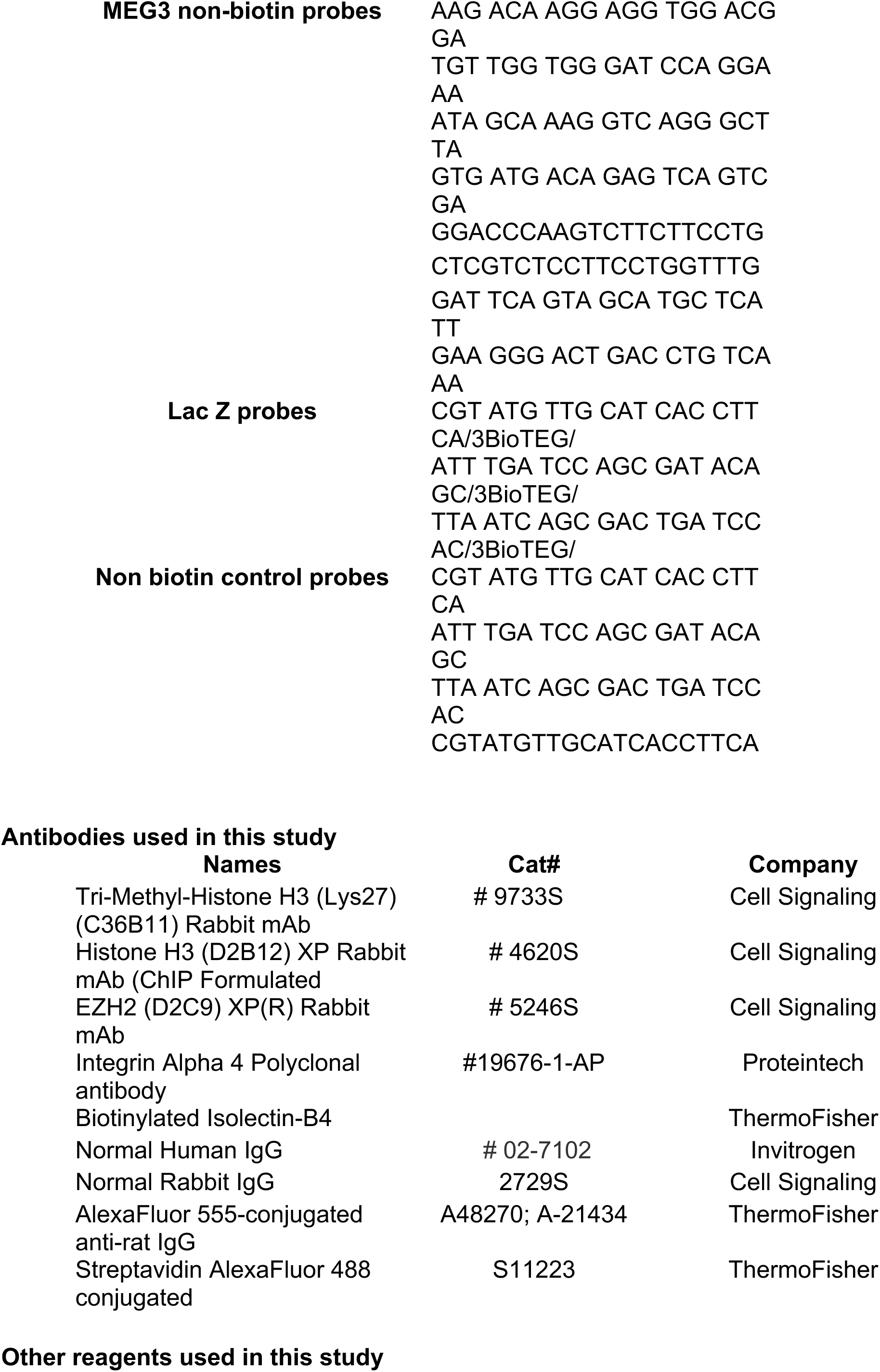
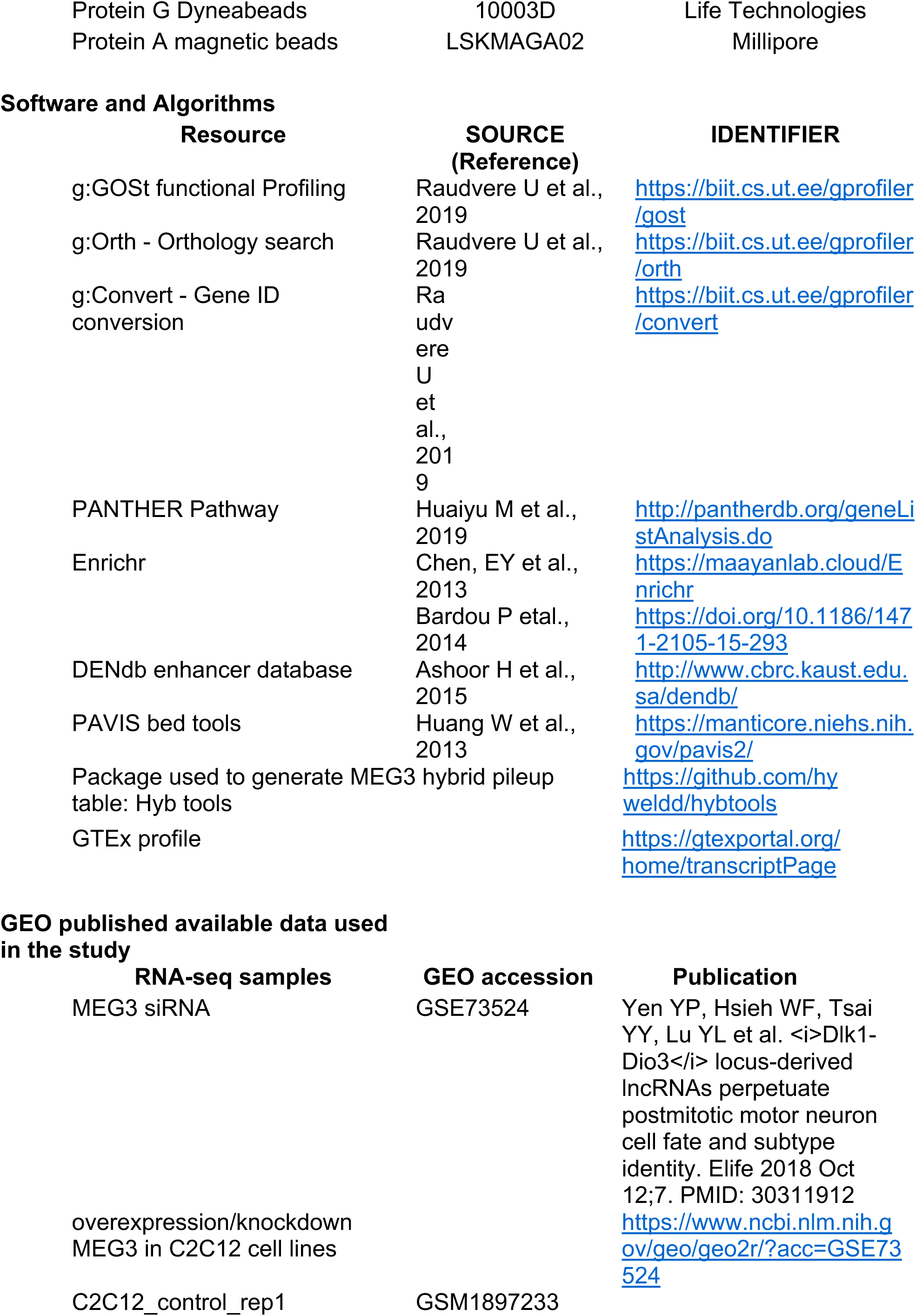
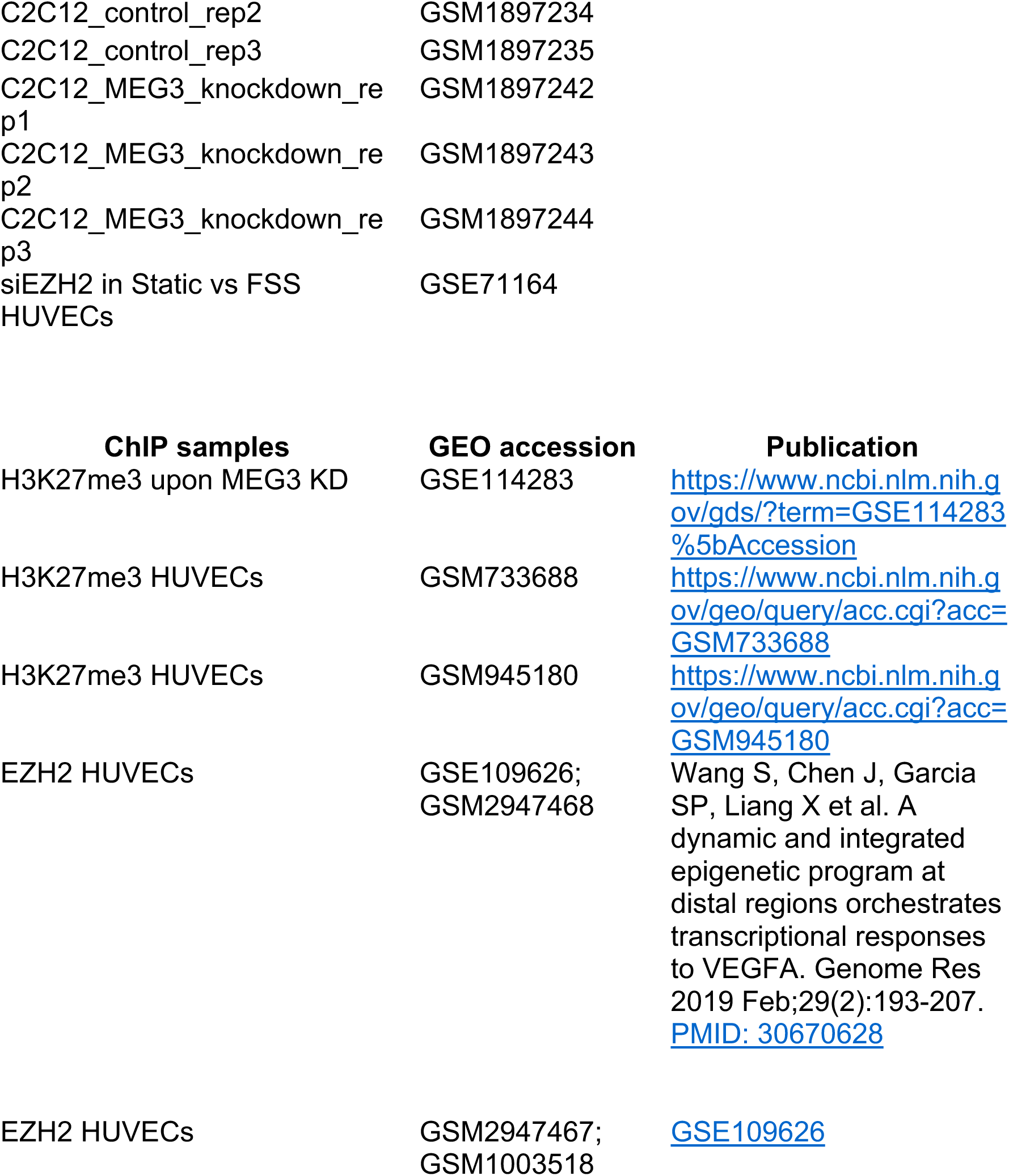
Primers, Antibodies and databases used in the study

**SUPPLEMENTARY FIGURE 1.**
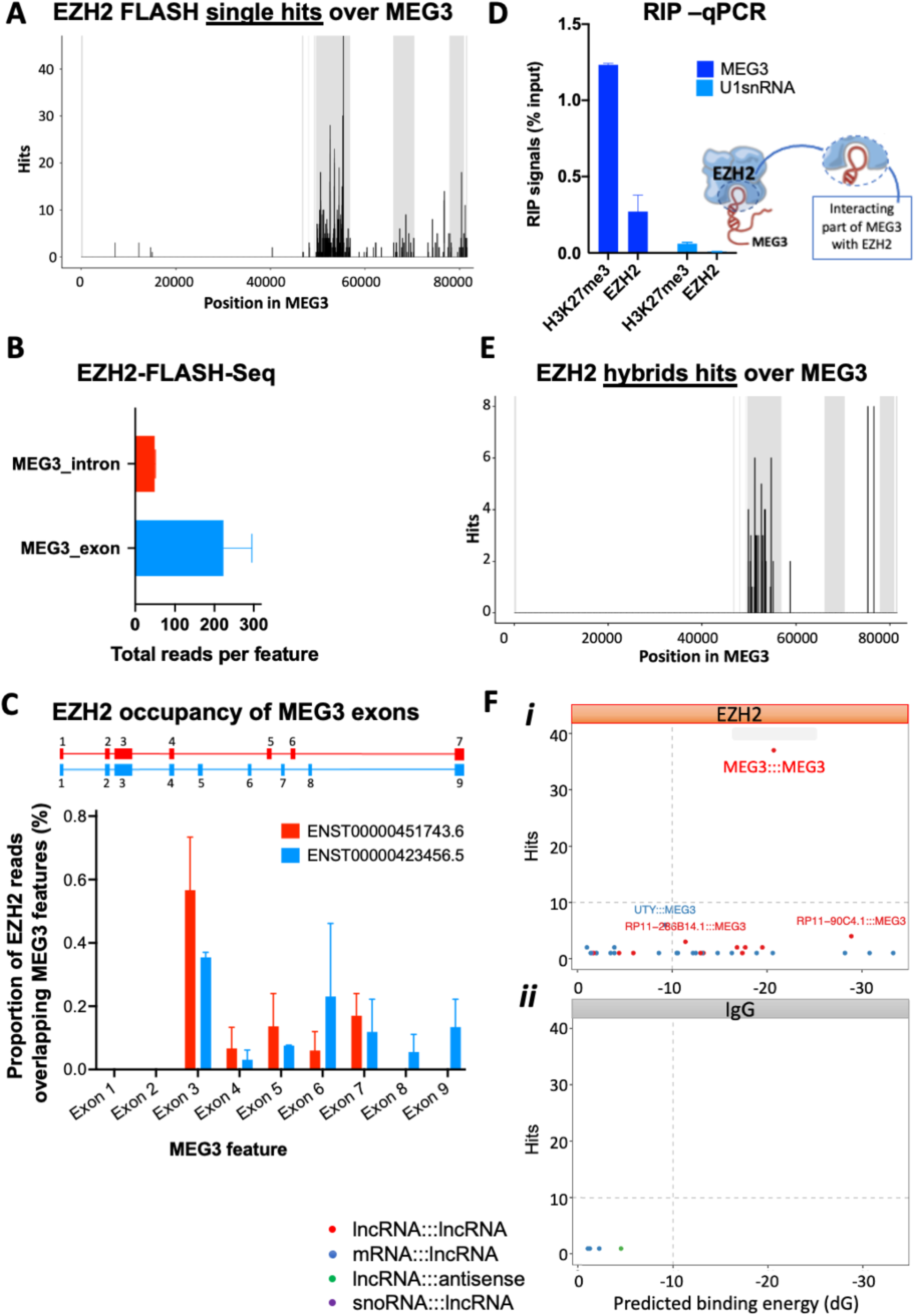
a) Distribution of annotated single hits over MEG3 gene, with statistically filtered EZH2-FLASH reads from two biological replicates in HUVECs. b) The occupancy of EZH2 hits over MEG3 features. Total reads per feature are given with exons being mostly occupies vs introns. c) Proportion of overlapping features over MEG3. The occupancy of EZH2 over each MEG3 exon is shown for two constitutively expressed transcripts. For both given transcripts there is high occupancy of exon 3. d) RNA immunoprecipitation (RIP) for EZH2 and H3K27me3 (repressive chromatin) followed by qPCR analysis. RIP-purified RNA from UV crosslinked HUVECs was used to prepare cDNA for qPCR analysis with primers against MEG3 (exon 3 region). Primers against U1snRNA gene serves as a negative control. Side diagram of EHZ2-MEG3 interacting region is charted as per FLASH hits and sequence. e) Distribution of EZH2 hybrids hits over MEG3 gene. Intermolecular MEG3-RNA interactions found in chimeras are captured by EZH2-FLASH-seq. Hits represent MEG3:MEG3 hybrids (black). IgG hybrids are plotted but are <1. f) Total MEG3:MEG3 hybrid count against predicted free energy of hybridization (dG) for MEG3 interactions (<u>red</u> lncRNA:MEG3, <u>blue</u> mRNA:MEG3, <u>green</u> MEG3:antisense, <u>purple</u> snoRNA:MEG3) with free hybridization energy cutoff at dG<-10 kcal mol^-1^, as captured by EZH2-FLASH-seq *(**i**) vs.* IgG control *(**ii**)*.

**SUPPLEMENTARY FIGURE 2.**
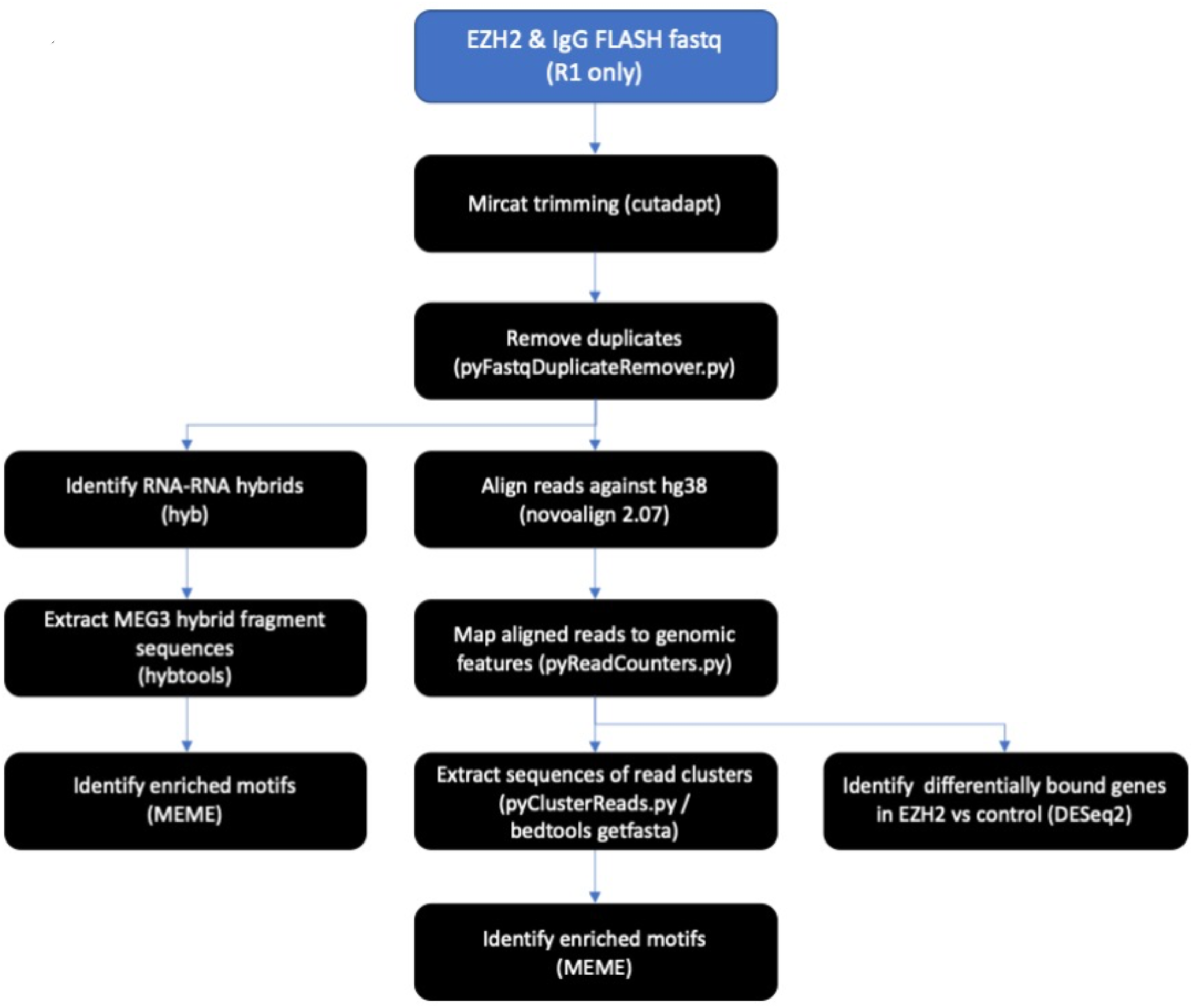
Computational analysis pipeline for FLASH-seq. Reads were aligned and used to identify EZH2-transcriptome. Differential expression (DE) analysis generated EZH2 bound genes vs. IgG control. Next, RNA:RNA-hybrids enriched in EZH2 RNA pull down were identified. The sequences of MEG3 hybrid fragments were specifically obtained and enriched motifs were identified.

**SUPPLEMENTARY FIGURE 3.**
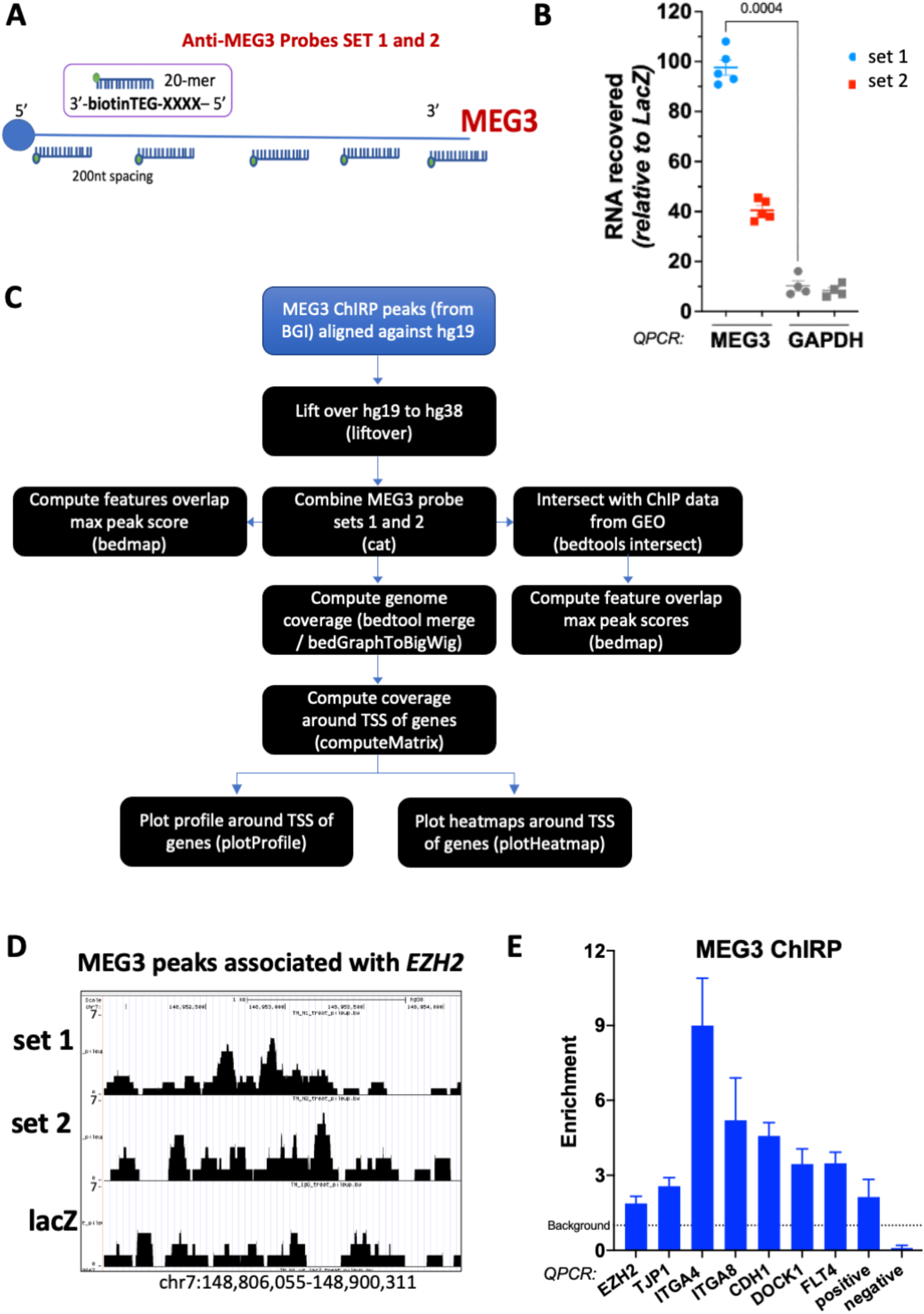
a) Overview of the design of probes against MEG3 gene that were divided in probe Set1 and Set 2. The biotynilated probes were of 20 nucleotides and were spaced out 200 nucleotides apart down the gene length. b) Validation of MEG3 probes specifically binding MEG3 gene, by ChIRP-qPCR in HUVECs. Pull down with probe set 1 or set 2 retrieved 100% and 40% RNA, respectively. GAPDH primers were used as control and MEG3-associated samples did not amplify. c) Computational analysis pipeline for ChIRP-seq outlining data processing. The peak coverage was within the 100bp window. d) MEG3-ChIRP peaks associated with *EZH2* gene as precipitated using both sets of probes (set 1 and 2). e) Enrichment of MEG3 signal by ChIRP-qpcr versus negative control (Background) at named promoter regions. MEG3 binding to genomic loci as validate by ChIRP-qPCR in HUVECs. Pull downs were performed with joined Odd and Even probes. Value 1 is a background level, defined by enrichment to LacZ negative probes in ChIRP. Control primers were designed for positive ChIRP peaks and used as a *positive control* and for regions deprived of MEG3-ChIRP reads as a *negative control*.

**SUPPLEMENTARY FIGURE 4.**
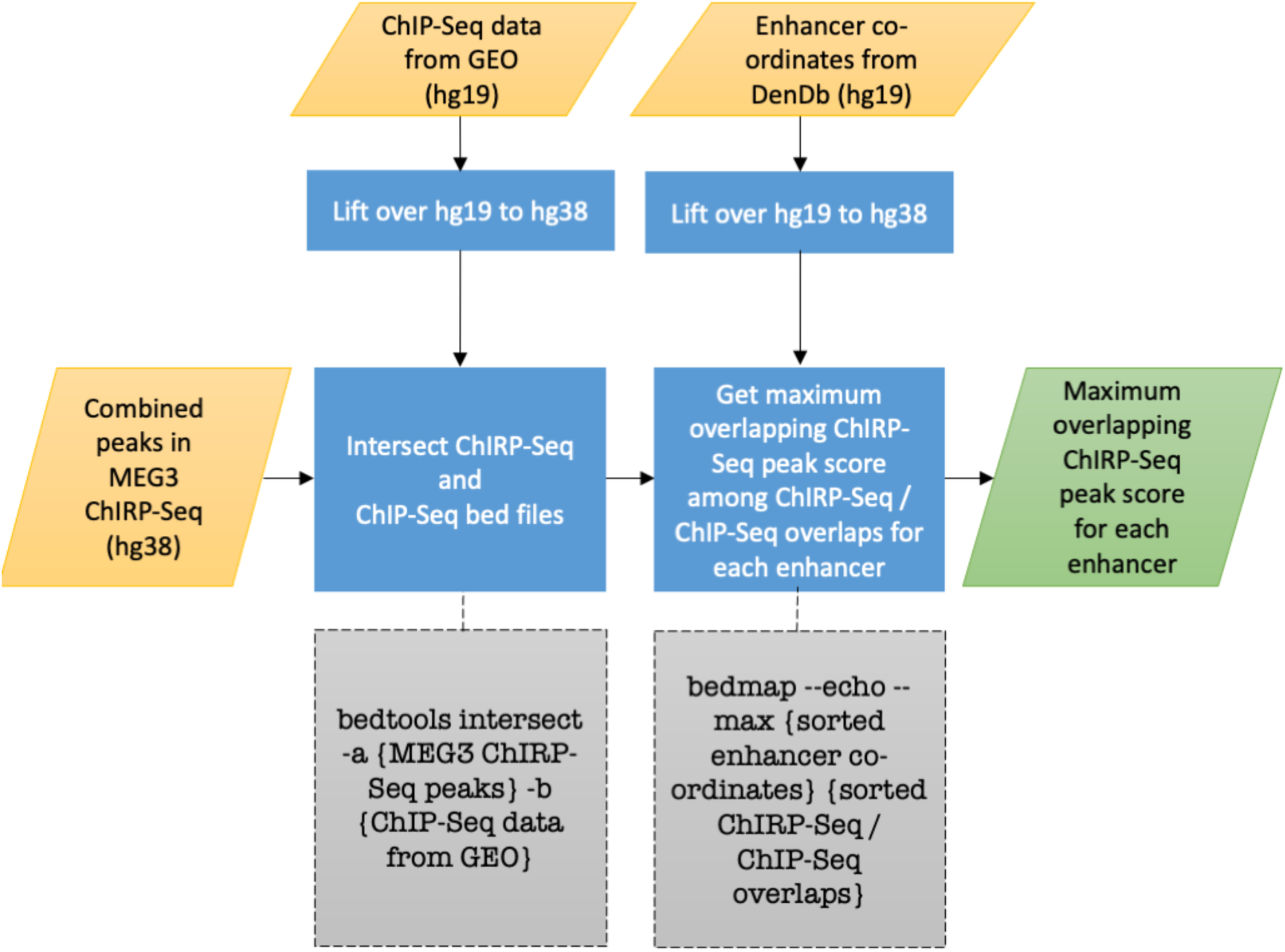
Computational analysis pipeline for ChIRP-seq intersection with ChIP-seq datasets. Enhancer coordinates were obtained from the DENdb database of integrated human enhancers.

**SUPPLEMENTARY FIGURE 5.**
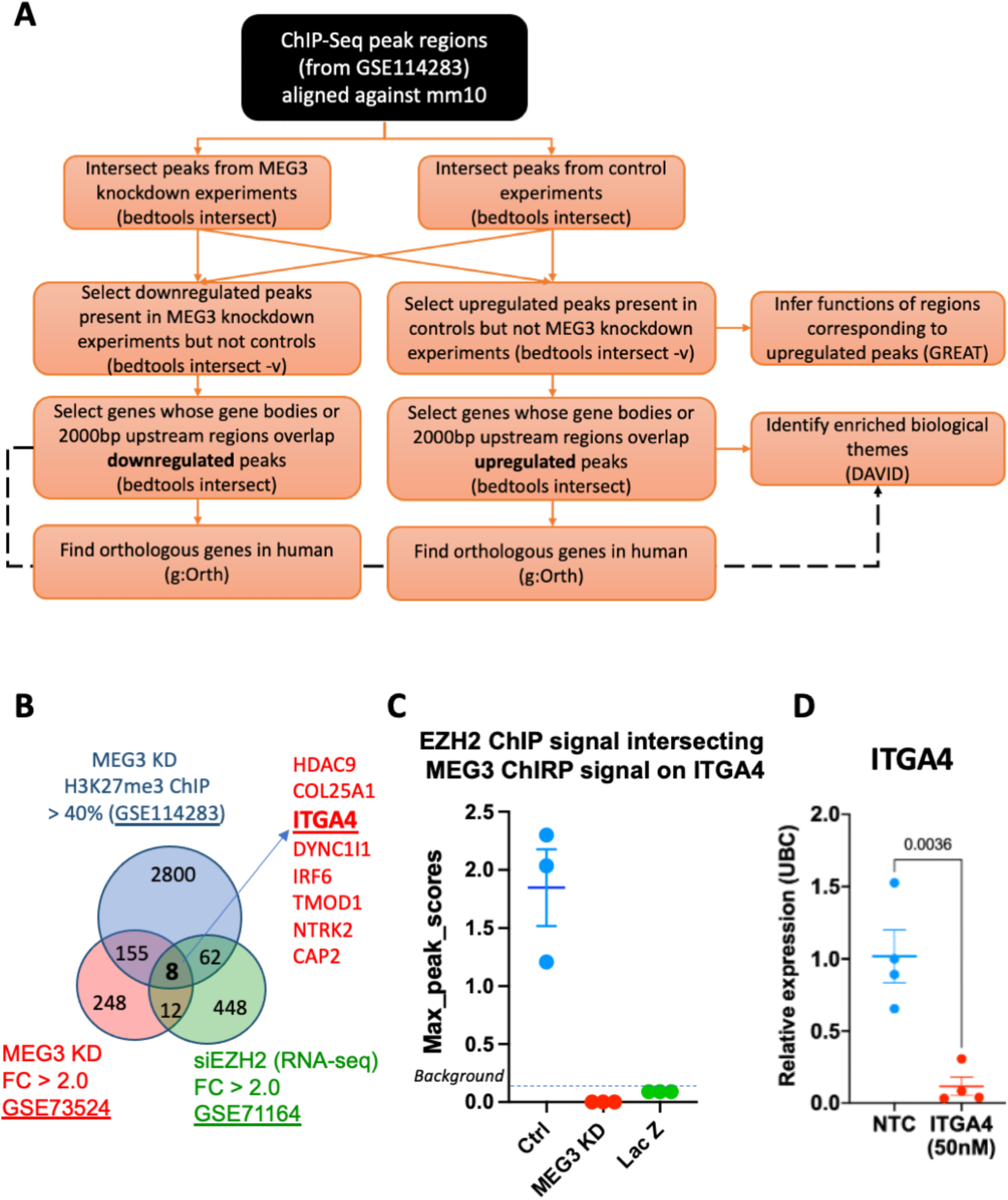
a) Computational analysis pipeline used to obtain orthologous peaks in human and intersect regions and genes enriched in repressive chromatin (H3K27me3) from ChIP-seq public dataset GSE114283. Up- and down-regulated genes were obtained associated with the peak region within 2000bp, and relevant function and biological pathway were associated using GREAT and DAVID analysis b) Overlap of the GEO datasets from *a* (Microarray **GSE73524**) and b (RNA-seq **GSE71164**) and the **GSE114283** ChIP-seq reads of H3K27me^3^ distribution in mouse MN cells depleted of MEG3 *vs.* control. ChIP extracted peaks unique to Ctrl *vs.* MEG3 KD were obtained, and associated mouse gene list composed based on reduction in H3K27me^3^ signal. Using gene orthologous analysis in gProfiler we obtained human orthologous targets that was used for data intersection. c) Maximum peak scores of the overlapping signal over ITGA4 promoter, obtained by intersection of EZH2 ChIP signal with MEG3-ChIRP signal at this region. Upon depletion of MEG3 the EZH2 signal is significantly reduced whereby no overlap with MEG3 ChIRP signal is seen. d) Relative expression of ITGA4 in HUVEC measuring the levels of ITGA4 following addition of siRNA (50nM).

**SUPPLEMENTARY FIGURE 6.**
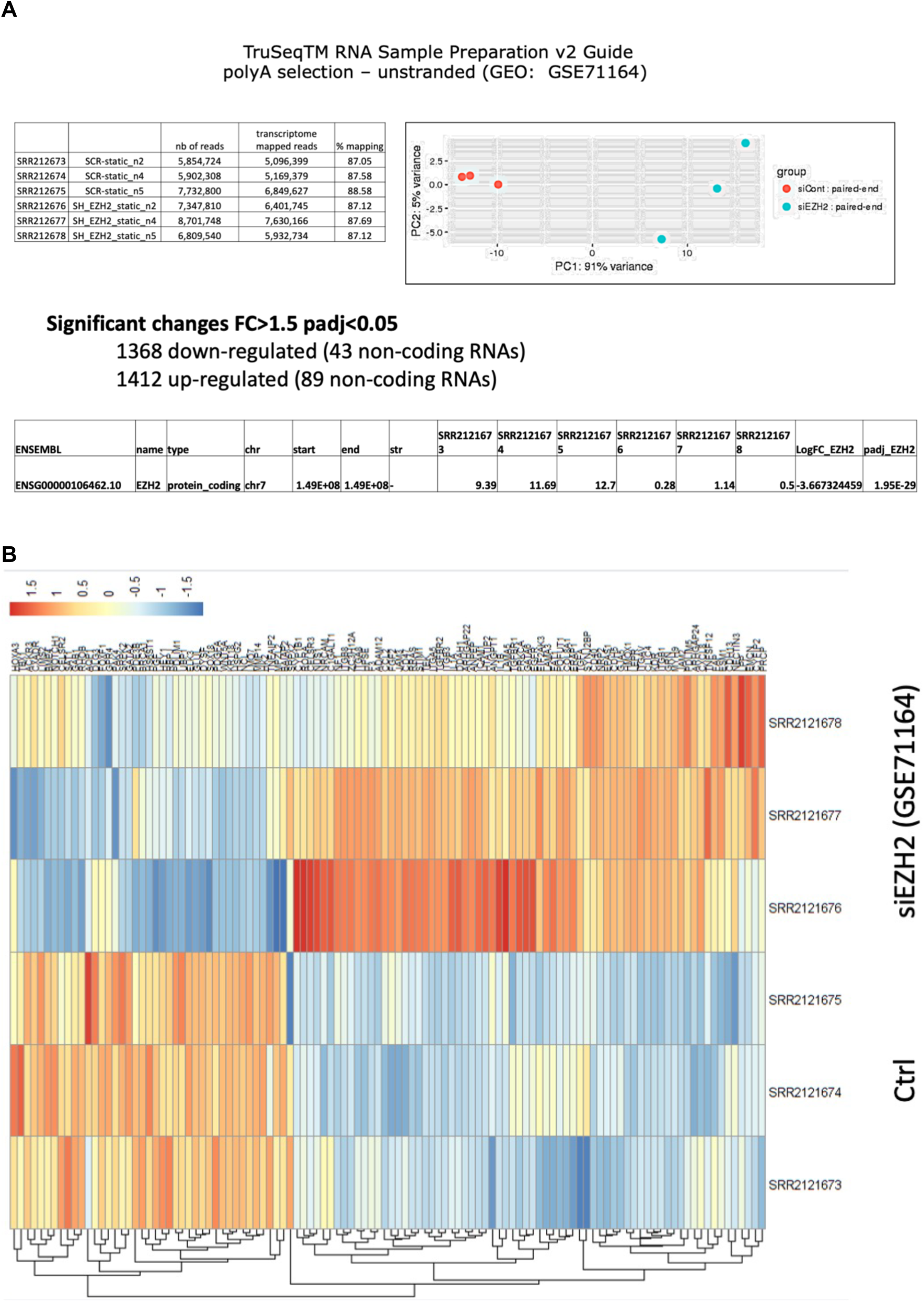
**a**) RNA-seq dataset from HUVEC cells depleted in EZH2 (GSE71164) was *de novo* analysed and mapped onto Hg38 with reads given in the table. The principal component analysis (PCA) was used to describe the variance between two groups (ctr *vs*. siEZH2); depletion of EZH2 gene is represented between samples (n=3) with reads per sample, in the bottom table. **b**) Heatmap of selected genes directly regulated by EZH2 and involved in angiogenesis and cell adhesion processes.

